# Structural visualization of the tubulin folding pathway directed by eukaryotic chaperonin TRiC

**DOI:** 10.1101/2022.03.25.483853

**Authors:** Daniel Gestaut, Yanyan Zhao, Junsun Park, Boxue Ma, Alexander Leitner, Miranda Collier, Grigore Pintilie, Soung-Hun Roh, Wah Chiu, Judith Frydman

## Abstract

**SUMMARY:** The ATP-dependent ring-shaped chaperonin TRiC/CCT is essential for cellular proteostasis. To uncover why some eukaryotic proteins can only fold with TRiC assistance, we reconstituted the folding of β-tubulin using human Prefoldin and TRiC. We find unstructured β-tubulin is delivered by Prefoldin to the open TRiC chamber followed by ATP-dependent chamber closure. CryoEM resolves four near-atomic resolution structures containing progressively folded β-tubulin intermediates within the closed TRiC chamber, culminating in native tubulin. This substrate folding pathway appears closely guided by site-specific interactions with conserved regions in the TRiC chamber. Initial electrostatic interactions between the TRiC interior wall and both the folded tubulin N-domain and its C-terminal E-hook tail establish the native substrate topology, thus enabling C-domain folding. Disordered CCT C-termini within the chamber promote subsequent folding of tubulin Core and Middle domains and GTP-binding. Thus, TRiC’s chamber provides chemical and topological directives that shape the folding landscape of its obligate substrates.

## INTRODUCTION

How proteins fold *in vivo* is a central unanswered question in biology. The growing list of diseases linked to protein misfolding, ranging from cancer to neurodegeneration, underscore the relevance of this question to human health (Buchner, 2019; Hartl et al., 2011; Labbadia and Morimoto, 2015; Mogk et al., 2018; Plate et al., 2016). Several classes of molecular chaperones play central roles in the biogenesis of newly translated polypeptides (Deuerling and Bukau, 2004; Deuerling et al., 2019; Willmund et al., 2013). While some chaperones, such as NAC or Hsp70 facilitate folding of a wide range of proteins, other, such as the ring-shaped chaperonin TRiC/CCT facilitate folding of a subset of proteins comprising ∼ 10% of the eukaryotic proteome (Yam et al., 2008). The proteins that require TRiC for folding, many of them essential for viability, are distinguished by their aggregation propensity and topological complexity (Stein et al., 2019; Yam et al., 2008). Interestingly, *in vivo*, TRiC functions in cooperation with upstream chaperones, including Hsp70 and the chaperone Prefoldin (PFD). Recent structural work showing PFD can bind directly to TRiC through an electrostatic pivot suggests a direct mechanism of substrate transfer (Gestaut et al., 2019). However, the underlying logic for the distinct *in vivo* requirement of specific proteins for TRiC and its cooperation with PFD, is not understood, and likely rooted in their sequence and folding requirements.

The contribution of TRiC/CCT to folding is particularly intriguing because this chaperonin assists folding of several eukaryotic proteins that cannot fold either spontaneously nor with any other chaperone, under any known condition (Balchin et al., 2018; Gao et al., 1993). While many different chaperones can bind proteins such as actin and tubulin and prevent their aggregation, only TRiC can promote their correct folding. An explanation for this obligate requirement is lacking, but perhaps the information present in their amino acid sequence does not suffice to reach the native state. In this hypothesis, TRiC may contribute the chemical and topological environment necessary to guide their folding, suggesting a co-evolution between TRiC and some eukaryotic proteins.

TRiC is a ∼1 MDa hetero-oligomer consisting of two identical rings stacked back-to-back (Cong et al., 2010). Each TRiC ring consists of eight unique paralogous subunits called CCT1-8 in a highly conserved arrangement (Leitner et al., 2012). Access to the central chamber of each ring is regulated in an ATP-dependent manner by a built-in lid, formed by flexible loops protruding from each subunit. The lid is open in the unliganded apo-state of TRiC, allowing substrates to bind through subunit specific contacts in the apical domains (Balchin et al., 2018; Joachimiak et al., 2014). ATP hydrolysis leads to lid closure, encapsulating the substrate within the central chamber, where folding occurs (Cong et al., 2012; Reissmann et al., 2012).

Unlike bacterial or archaeal chaperonins, TRiC is assembled from eight different subunits in a conserved arrangement (Leitner et al., 2012). The hetero-oligomeric nature of TRiC is thought to be key to its unique ability to fold obligate substrates. The specific subunit arrangement segregates substrate and ATP binding properties within the ring and produces an asymmetric charge distribution within the closed chamber (Leitner et al., 2012; Reissmann et al., 2012). Supporting a role for subunit divergence contributing to the folding process, previous studies showed unfolded substrates such as actin, AML or VHL engage open TRiC through subunit specific contacts with CCT apical domains (Balchin et al., 2018; Roh et al., 2016; Spiess et al., 2006; Tam et al., 2009). This could bias the initial topology of the TRiC-bound substrate to a conformation that is productive for folding. Analysis of TRiC-mediated actin folding indicates that the gradient of ATP affinities within the ring promotes stepwise release during chamber closure to facilitate folding (Balchin et al., 2018; Reissmann et al., 2012). However, how folding occurs inside the closed TRiC folding chamber is not understood. Is the bound substrate released into the ATP-closed chaperonin chamber, as shown for bacterial chaperonin GroEL-ES? How do the properties of the TRiC closed chamber affect the folding pathway of its substrates?

To address these fundamental questions, we reconstituted the chaperone mediated folding pathway of the archetype TRiC obligate substrates; α- and β-tubulin. Since, in the cell, TRiC cooperates upstream with another hetero-oligomeric chaperone Prefoldin (PFD) for substrate delivery, we initiated the folding reaction starting with purified PFD-tubulin complex. We find PFD maintains tubulin in a highly unstructured state and directly transfers substrate into the internal TRiC chamber. ATP addition then leads to tubulin encapsulation and folding within the closed TRiC chamber. A combination of biochemical and cryogenic electron microscopy (cryoEM) structural analyses of each step in the chaperone-mediated folding reaction strikingly reveals specific contacts with the inner chamber of TRiC direct the tubulin folding trajectory. Electrostatic and polar contacts with individual CCT subunits promotes domain-wise formation of four progressively folded tubulin intermediates culminating in the native state.

## RESULTS

### Reconstituting the human Prefoldin-TRiC pathway for efficient tubulin folding

Tubulin is folded *in vivo* by a chaperone pathway consisting of PFD and TRiC/CCT (Geissler et al., 1998; Vainberg et al., 1998). α or βTub monomers folded by TRiC subsequently bind to assembly factors TBCB or TBCA respectively, which steer the folded monomers through a network of factors that assemble tubulin dimers (Lewis et al., 1997; Nithianantham et al., 2015). We reconstituted the PFD-TRiC-tubulin monomer folding pathway using purified human components (Figure 1A, S1A-C). We generated human PFD-bound to either α or βTub by co-expression in insect cells (Figure S1C) and incubated them with purified human TRiC in the presence and absence of ATP. Folding reactions were analyzed by native polyacrylamide gel electrophoresis (N-PAGE) and limited-proteolysis followed by gel electrophoresis (LP-PAGE) (Figure 1B, C). N-PAGE analyses of these reactions indicated that, upon TRiC addition, the initial PFD-αTub or PFD-βTub complexes formed a ternary complex with TRiC, reflected in a characteristic mobility shift (Figure 1B for βTub and S1D for αTub). Subsequent addition of ATP led to production of folded tubulin monomer, reflected by formation of protease-resistant tubulin as well as formation of a tubulin complex with assembly factors TBCB or TBCA (Figure 1B; S1D).

**Figure 1:**
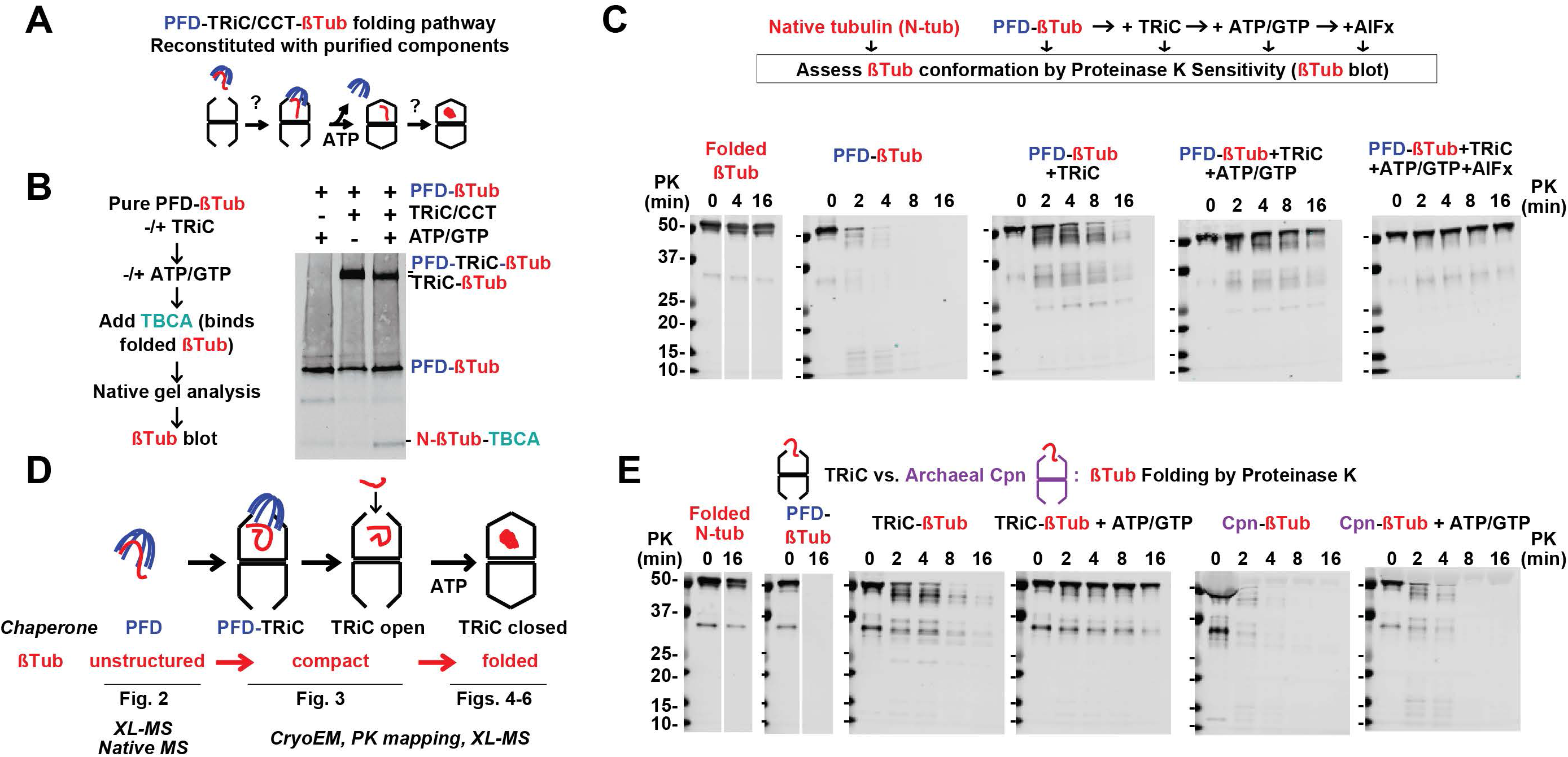
Reconstitution of the Tubulin folding pathway from purified components. (A) Schematic of Tubulin folding pathway reconstituted from purified components to assess in vitro how TRiC assists Tubulin to reach its native fold. (B) Validation of recombinant system; βTub co-expressed/purified with PFD comigrates on a native gel. In the absence of nucleotide βTub comigrates with TRiC, and addition of nucleotide (ATP/GTP) drives folding and partial release of βTub in the presence of TBCA. (C) Protease sensitivity assay (Proteinase K), establishes Tubulin bound to PFD is protease sensitive, transfer to TRiC from PFD causes a decrease in sensitivity, and addition of nucleotide drives folding resulting in limited protease sensitivity equivalent to Tubulin in a native α/β Tubulin dimer. Locking TRiC in a closed conformation with an ATP-AlFx also provides protection from protease. (D) After validation we studied each step of the folding pathway using the indicated techniques including XL-MS, Native-MS, CryoEM, and PK mapping. (E) PK assay demonstrating βTub copurifed with TRiC is folded upon ATP/GTP addition, while the simpler homologous group II chaperonin CPN found in Archaea is unable to provide protease protection upon ATP/GTP addition.

We next characterized the conformation of βTub along the PFD-TRiC chaperone pathway using limited proteolysis, which tests the sensitivity of folding intermediates to Proteinase K (LP-PAGE, Figure 1C, S1E). Similar results were obtained for βTub and αTub suggesting a shared mechanism of chaperone action for these proteins (Figure S1D, E). As expected, folded tubulin in αβ dimers is nearly fully protease resistant, except for some degradation of flexible C-terminal tails. In contrast, both αTub and βTub bound to PFD were highly protease sensitive, indicative of an unfolded dynamic conformation with flexible regions susceptible to rapid Proteinase K digestion. Addition of TRiC and ensuing formation of a ternary PFD-tubulin-TRiC complex, led to enhanced αTub and βTub protection against Proteinase K. We obtained similar results using an antibody directed against the N-terminal domain of tubulin as well as one directed against the C-terminus of tubulin (Figure S1G-I). These experiments indicate tubulin becomes compacted upon binding to TRiC. Interestingly, probing with the N-terminal directed antibody but not the C-terminal directed antibody also detected smaller protected protease fragments, suggesting formation of partially folded N-terminal domain structures. Incubation of the PFD-tubulin-TRiC complex with ATP, which promotes TRiC chamber closure and PFD dissociation (Gestaut et al., 2019b), led to formation of full length protease protected tubulin. The level of protection was similar to that of folded αTub and βTub in the native assembled tubulin dimer. Since under these ATP conditions TRiC cycles between open and closed states, these experiments suggest tubulin is folded by TRiC in an ATP-dependent manner. To confirm this, TRiC-tubulin was incubated with ATP to promote folding, followed by subsequent addition of apyrase to hydrolyze all ATP causing the TRiC lid to reopen. Under these conditions, tubulin remained fully protease protected, confirming the ATP-induced protease-protection arises from tubulin reaching the folded conformation (Figure S1G, I). Of note, locking the TRiC chamber into a closed conformation by the addition of ATP-AlFx, also led to strong protection from proteolysis. This indicates the TRiC-bound substrate is fully encapsulated within the central chamber.

These experiments suggest PFD and TRiC differentially affect the conformation of the tubulin polypeptide (Figure 1D). PFD maintains tubulin in an unstructured and highly protease sensitive state, with no obvious protease-protected fragments observed. Upon PFD transfer of tubulin to TRiC, the substrate becomes compacted, acquiring some protease-resistance. Finally, ATP-driven encapsulation leads to a fully protease-resistant tubulin that can bind to the assembly factors recognizing folded monomers. We next examined tubulin folding when the PFD-delivery step is bypassed and tubulin is directly bound to TRiC (Figure 1E). Limited proteolysis analysis indicated tubulin adopts the same compact intermediate upon TRiC binding and also reaches a fully protease protected conformation upon ATP/GTP addition (Figure 1E). Importantly, when PFD-βTub is delivered to the TRiC-like archaeal chaperonin from *M. maripaludis*, Mm-Cpn, it does not achieve full protease protection upon ATP/GTP incubation (Figure 1E). Thus, TRiC is primarily responsible for tubulin folding, while PFD functions to maintain tubulin in an unstructured state, and prevent formation of off-pathway intermediates. Having reconstituted PFD-TRiC mediated tubulin folding, we set out to study each stage of the pathway using a hybrid structural and biochemical approach involving crosslinking, mass spectrometry and cryoEM (Figure 1D).

### Defining PFD interactions with non-native tubulin

PFD is a jellyfish-shaped complex with six different subunits forming a double beta-sheet barrel from which coiled-coils extend forming “tentacles” (Figure 2A). The flexible nature and relatively small size of the PFD-Tub complex hinders its structural characterization by cryoEM. Accordingly, we mapped the PFD-Tub contacts using cross-linking-mass spectrometry (XL-MS), with similar results obtained for PFD-bound αTub and βTub (Figure 2A, B; Figure S2A). Within PFD, the observed substrate XL mapped primarily to the coiled coil tentacles and their disordered tails, indicating non-native tubulin binds inside the PFD “chamber” (Figure 2B; S2B). Positional information on the PFD-Tub contacts within the chamber, visualized by mapping contacts between the tubulin polypeptide and PFD revealed binding occurs in a highly dynamic interaction (Figure 2C, D). The N-terminal domain of tubulin made numerous contacts throughout the PFD inner chamber, with most crosslinks (XL) mapping to contiguous PFD1/5/6 coiled coils. In contrast, the tubulin core domain was crosslinked primarily to one specific region in PFD6. The C-terminal domain of tubulin also crosslinked to a continuous patch on PFD3/5/6, while the M domain crosslinked to several points at the bottom of the PFD chamber. These analyses suggest tubulin adopts a highly dynamic conformation, consistent with the high protease sensitivity observed for PFD-bound tubulin, even though some regions of tubulin bind more stably to specific regions within the PFD chamber.

**Figure 2:**
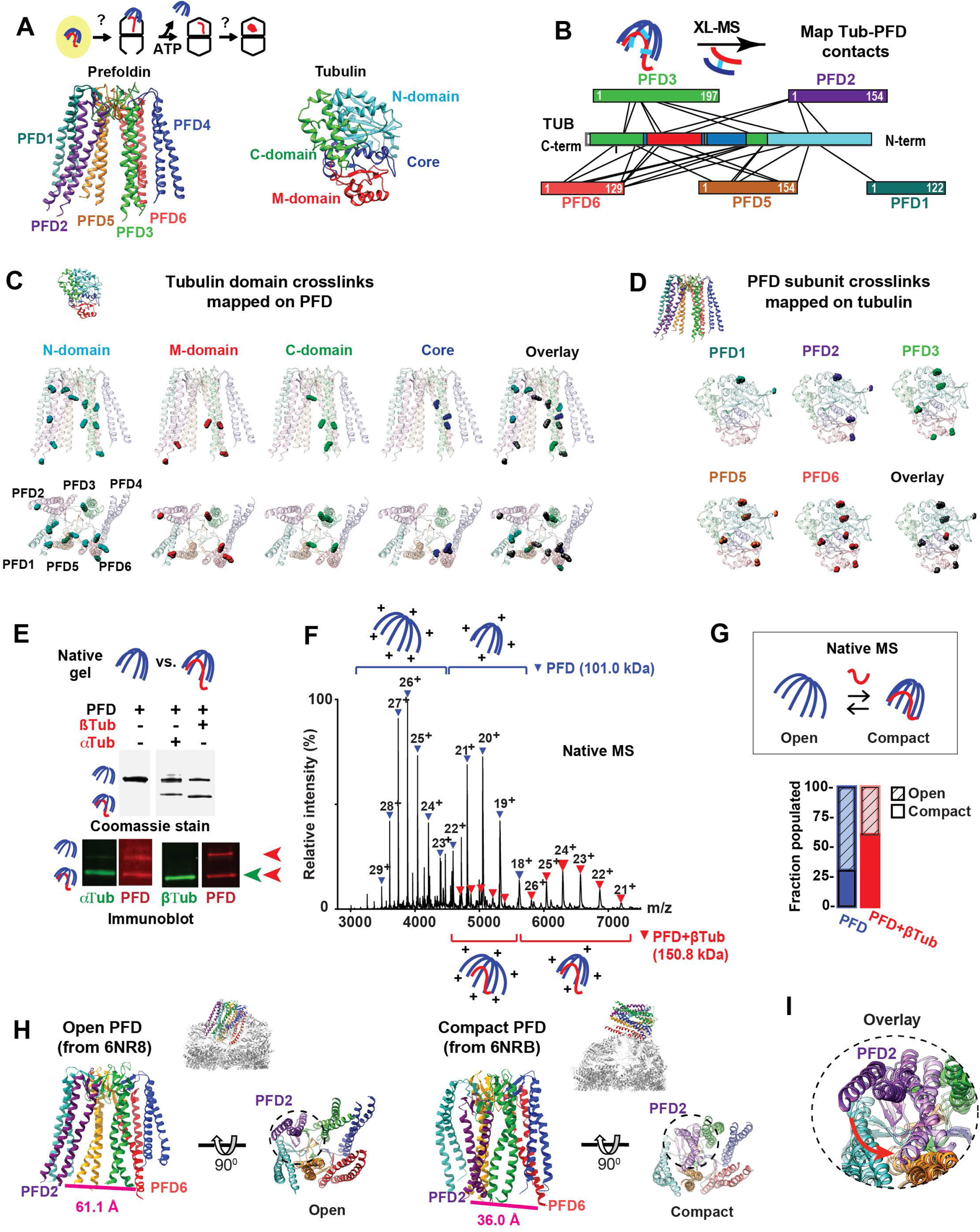
Defining PFD interactions with non-native tubulin. (A) Schematic model of Tubulin folding pathway with PFD-Tubulin highlighted and PFD and Tubulin structures with color scheme for PFD subunits and Tubulin domains. (B) Crosslinking Mass spectrometry (XL-MS) results from PFD-αTub and PFD-βTub mapped on to linear bar diagrams of subunits, Tubulin is colored by domain. (C) Site of XL between Tubulin domains and PFD mapped on to PFD structure with overlay of all XL, note XL to unstructured regions of PFD tentacles are mapped on to the last resolved residue. (D) XL between each subunit of PFD and Tubulin mapped on to Tubulin structure with overlay of all XLs. (E) Native gel showing PFD migrates as a single band in the absence of substrate, while a faster migrating band is produced when copurified with α or βTub. Western blot establishes the faster migrating band is PFD bound to Tubulin. (F) Representative native mass spectrum of PFD mixed with βTub. Two charge state distributions are observed for both PFD and the PFD-βTub complex. These can be inferred to correspond to different conformations in solution which we term “open” or “compact”, represented by cartoons with greater or lesser surface area and thus more or fewer charges (+). (G) Quantification of the fractions of PFD and PFD-βTub populating open and compact conformations by native mass spectrometry. Upon binding βTub, the PFD conformational equilibrium shifts to favor the compact form. Because buffer exchange and ionization conditions could affect conformations, these data constitute a comparison within the same mass spectra and should not be considered absolute fractions. (H) Structures of PFD from the least engaged (PDB:6NRB) and most engaged (PDB:6NR8) states with TRiC with the distances between the distance the terminal residue of PFD2 (Ile 124) and PFD6 (Glu 114) indicated. (I) Overlay of PFD structures focusing on PFD2 (purple) movement in least engaged (light color) and most engaged (dark color) states

PFD is an ATP-independent chaperone, raising the question of how substrate binding and release are regulated. N-PAGE of PFD αTub and βTub-complexes suggested substrate binding induces a conformational compaction of PFD, since PFD-substrate complexes migrate faster than PFD alone (Figure 2E). This observation was further substantiated by native mass spectrometry analyses (Figure 2F, G, S2C, D). PFD alone produced a bimodal charge distribution for the single expected PFD mass (101 kDa) (Figure S2C, D). Because the charge states produced during gentle ionization are strongly dependent on the exposed surface area (Collier et al., 2019; Kaltashov and Mohimen, 2005; Santambrogio et al., 2012; Scott et al., 2015), these data indicate pure PFD adopts two distinct structural states: one more compact and one more extended. Analysis of PFD-βTub complexes by native mass spectrometry also yielded a bimodal charge series consistent with the expected mass of the binary PFD-βTub complex (151 kDa) (Figure 2F, S2E). Of note, the relative intensity, and therefore abundance, of the compact species was higher in PFD-βTub peaks compared to PFD peaks (Figure 2E-G, S2E). This indicates, alongside N-PAGE, that tubulin binding induces a shift to the more compact PFD conformation. The extensive XL between tubulin and the tentacles of PFD2 and PFD6, located at opposite sides of the PFD chamber, suggest a rationale for this compaction, while previous structural analyses of the PFD-TRiC interaction suggest a possible function (Figure 2H, S2B, F, G). Thus, previous CryoEM of the empty PFD-TRiC binary complex found a flexible contact point between the base of PFD and the apical domain of CCT4 that allows the PFD tentacles to pivot into the chamber (Gestaut et al., 2019). While PFD adopts a compact conformation in the initial TRiC contact but shifts to an extended conformation when the PFD-TRiC chambers become aligned (Figure 2H, I; S2F, G). This suggests PFD initially binds TRiC in the compact, substrate-bound conformation but then chaperonin interactions promote the extended PFD state to facilitate substrate release into the TRiC chamber.

### The ternary complex of PFD-tubulin-TRiC in the open conformation

To visualize the PFD to TRiC substrate handoff, we added PFD-βTub to purified TRiC and studied the ternary complex by cryoEM (Figure 3A; S3A-D). Our analysis of the PFD-βTub-TRiC ternary complex showed the chamber of PFD fully aligned with that of TRiC through subunit specific contacts (Figure 3B). Strikingly, we observed a significant extra density inside the inter-ring space formed at the TRiC equatorial region, that is not observed in the absence of substrate (Figure 3B-D) (Gestaut et al., 2019). The PFD tentacles extend into the TRiC chamber, with the PFD6 coiled-coil strikingly making a clear connection to the extra density in the inter-ring space (Figure 3C). The PFD in this TRiC complex is in the open extended conformation, suggesting TRiC binding reduces its affinity for the bound substrate. Indeed, no significant density is observed inside the PFD chamber, despite the substantial extra density observed between the two TRiC rings. The oval-shaped substrate-induced extra-density is asymmetrically located within the inter-ring TRiC chamber, close to subunits CCT8, 6, and 3. Of note, we also observe clear connections between this density and the N-terminal tails of CCT5 and 7, which are normally not resolved in open TRiC structures (Figure 3D).

**Figure 3:**
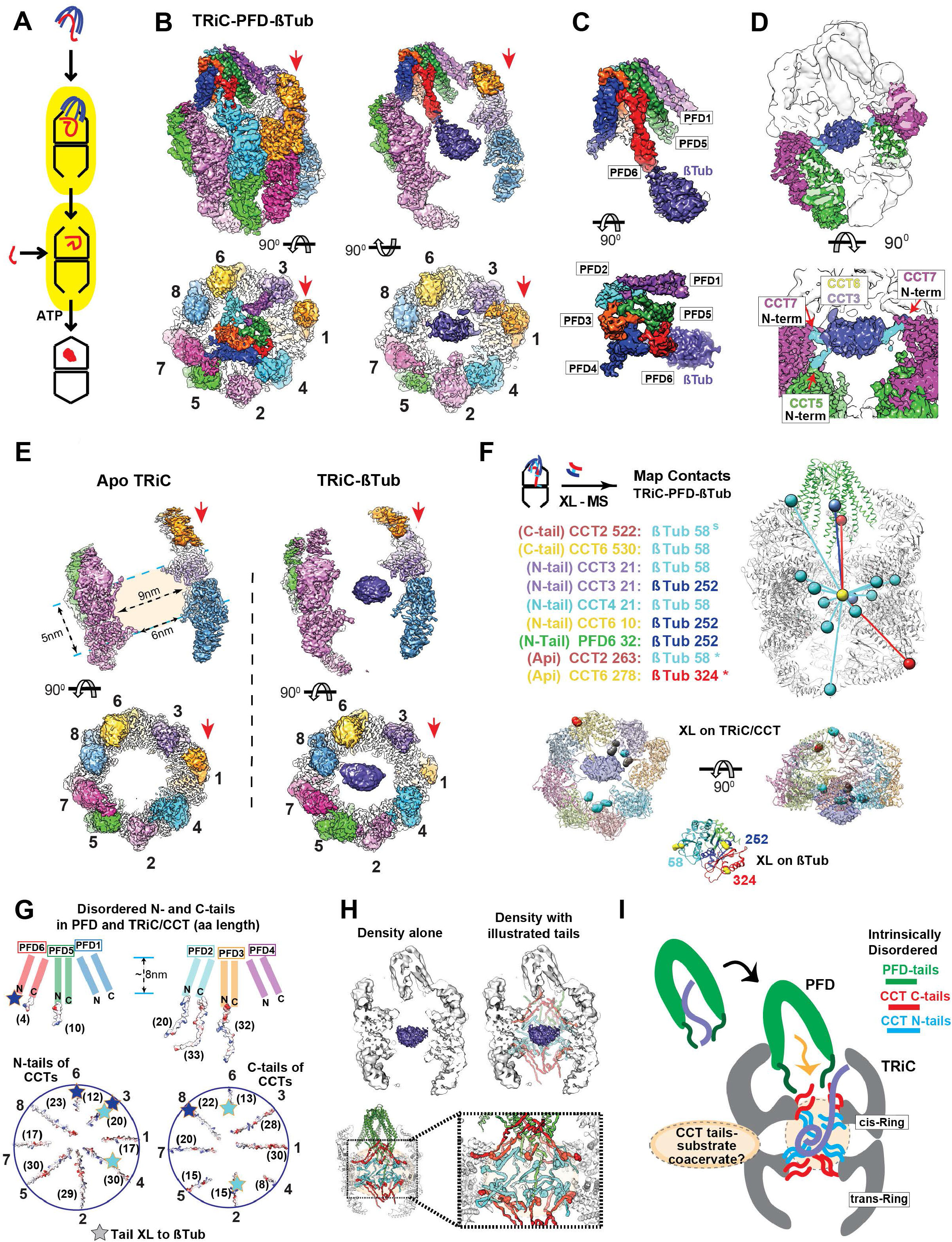
The ternary complex of PFD-tubulin and TRiC in the open conformation. (A) Schematic model of substrate loading to TRiC. (B-D) 3D-reconstruction of TRiC- PFD-βTub ternary complex, (B) Side and end-on view of the ternary complex, and slice views showing βTub density bound to TRiC. Red arrows indicate CCT1. (C) Side and end-on views segmented PFD-βTub density. (D) Side and top view at lower threshold to illustrate N-terminal tail contacts from CCT5 and CCT7. (E) Side and top views from 3D- reconstruction of apo-TRiC and TRiC-βTub highlighting the βTub density is absent from apo-TRiC. Red arrows indicate CCT1. (F) TRiC-βTub inter-molecular XL mapped onto the model, note XL are mapped to the center of mass of the density and colored by the domain of Tubulin they XL to. Residues that XL to βTub are also highlighted on a single ring of TRiC and colored by the domain of Tubulin they XL to, residues that XL to multiple domains are colored grey. XL between βTub and TRiC apical domains (*) are from (Joachimiak et al., 2014) with denatured Tubulin, and XL denoted with a (^s^) came from (Knowlton et al., 2021). (G) Schematic and electrostatic surface illustration of disordered tail residues from N and C termini of PFD and TRiC. Star mark in blue indicates the location of the connected density form PFD to βTub in the ternary complex.(H) Schematic modeling of tails into the TRiC inter-chamber space with map and model.(I) Cartoon illustration of substrate delivery by the interplay of tails between TRiC and PFD. The dotted circle in the TRiC chamber represents hypothetical coacervate formed by substrate polypeptide and disordered CCT tails.

We next examined whether the extra density indeed arises from βTub. We prepared a TRiC-βTub complex generated without PFD and carried out cryoEM analyses of both substrate-free apo-TRiC and TRiC-βTub. Only the βTub-containing TRiC has density in the central inter-ring region, while no notable density is observed in substrate-free apo-TRiC (Figure 3E). Extensive focused 3D classification on the extra-density failed to discern any folded domain or secondary structure within the density. While the presence of substrate-dependent density indicates the formation of a compact state, this structural uncertainty suggests heterogeneity in the folding state and positioning of βTub in the open conformation of TRiC. Our experiments indicate that in the open conformation of TRiC, there is substrate density in a chamber between the rings created by the equatorial domains of CCTs protruding into the central axis, which form ∼6 nm open cap covering a ∼9×5 nm globular inner space (Figure 3E).

Cross-linking mass spectrometry (XL-MS) analyses further characterized the contacts of βTub with PFD and TRiC within the ternary complex and in the binary βTub-TRiC complex (Figure 3E; S2A and Table S2). Notably, TRiC addition to PFD-βTub led to a near complete loss of βTub’s XL to PFD and appearance of extensive XL to TRiC, consistent with substrate transfer between these chaperones (Figure 3F). The termary complex only had one βTub XL to PFD1 and one βTub XL to the N-terminus of PFD6, confirming the proximity of the tip of the PFD6 tentacle to the substrate density (Figure 3E). In addition, we observed multiple PFD6 XL to the tails of CCT1 and CCT2 in the equatorial region of TRiC (Figure S3E, F). Importantly, the XL-MS analyses indicated extensive contacts between the substrate and the N and C terminal tails extended from the equatorial domains of CCT subunits, confirming the location of the βTub density in cryoEM maps (Figure 3F, G XL between βTub and the CCT apical domains were observed in a TRiC-βTub complex generated without PFD but not detected in either the PFD-βTub-TRiC ternary sample (Cuellar et al., 2019; Joachimiak et al., 2014). Similarly, we did not observe notable substrate density at the similar contour level in the apical domain regions of TRiC. Perhaps the substrate interactions with apical domains are too dynamic to be clearly observed, or alternatively apical contacts serve in the initial capture of substrate for subsequent engagement with the CCT tails. The ternary complex with PFD may thus reflect the substrate state after transfer to the central chamber.

Historically, TRiC has been considered as a two-chamber folding system, but our data suggests the existence of a third chamber that holds the substrate in the open TRiC conformation. While theoretically, this third chamber provides enough space for ∼50 kDa protein, it is important to note that the unresolved intrinsically disordered N- and C-termini tails of CCTs projecting into this space comprise ∼35.5 kDa of protein mass (Figure 3G). Our observations of substrate density by cryoEM and increased protection from proteinase K indicate the substrate becomes compacted upon TRiC binding, yet we could not recognize any stable domain formation. Remarkably, the unstructured CCT tails filling this space and contacting βTub in the open state contain both polar as well as hydrophobic character (Figure 3G-H). We speculate that these disordered tails may function as a tethered solvent that maintains the unfolded polypeptide in a conformationally dynamic coacervate state, thus preventing formation of trapped intermediates. In this manner, tubulin folding intermediates may be stabilized by interactions with CCT tails in the open chaperonin conformation (Figure 3H, I).

### CryoEM identifies βTub folding intermediates within the ATP-closed TRiC chamber

The next step of the βTub folding cycle occurs upon closure of TRiC with ATP hydrolysis (Figure 4A). We incubated the TRiC-βTub complex with ATP-AlFx to stabilize the closed conformation and examined βTub folding in the chamber by cryoEM. While the 3.8 Å open TRiC state contains the βTub density in the inter-ring space (Figure 3; S4A, B), the ATP-closed TRiC conformation contains the substrate density in the apical region within one of the chambers (Figure 4A). The density map of closed TRiC at 2.8 Å resolution revealed the repositioned βTub had well-defined secondary structural features (Figure 4A). Additionally, most of the CCT N- and C-terminal tails could be traced in the closed conformation but not in the open conformation. The initial overall map of closed TRiC-βTub visualized partially folded βTub with varying resolution suggesting compositional and conformational heterogeneity of βTub. We therefore performed 3D classification focusing on the βTub density, which identified four distinct conformations of βTub within the TRiC chamber, which were reconstructed at resolutions ranging from 2.9 Å to 3.6 Å (Figure 4B, S4C-E). The quality of the maps allowed us to build models of each TRiC bound folding intermediate (Figure 4C). These four maps display densities connecting secondary structure elements, including α-helices and β-sheets, are suggestive of four progressively folded βTub intermediates in the closed TRiC chamber (Figure 4B,C; Movie S1).

**Figure 4:**
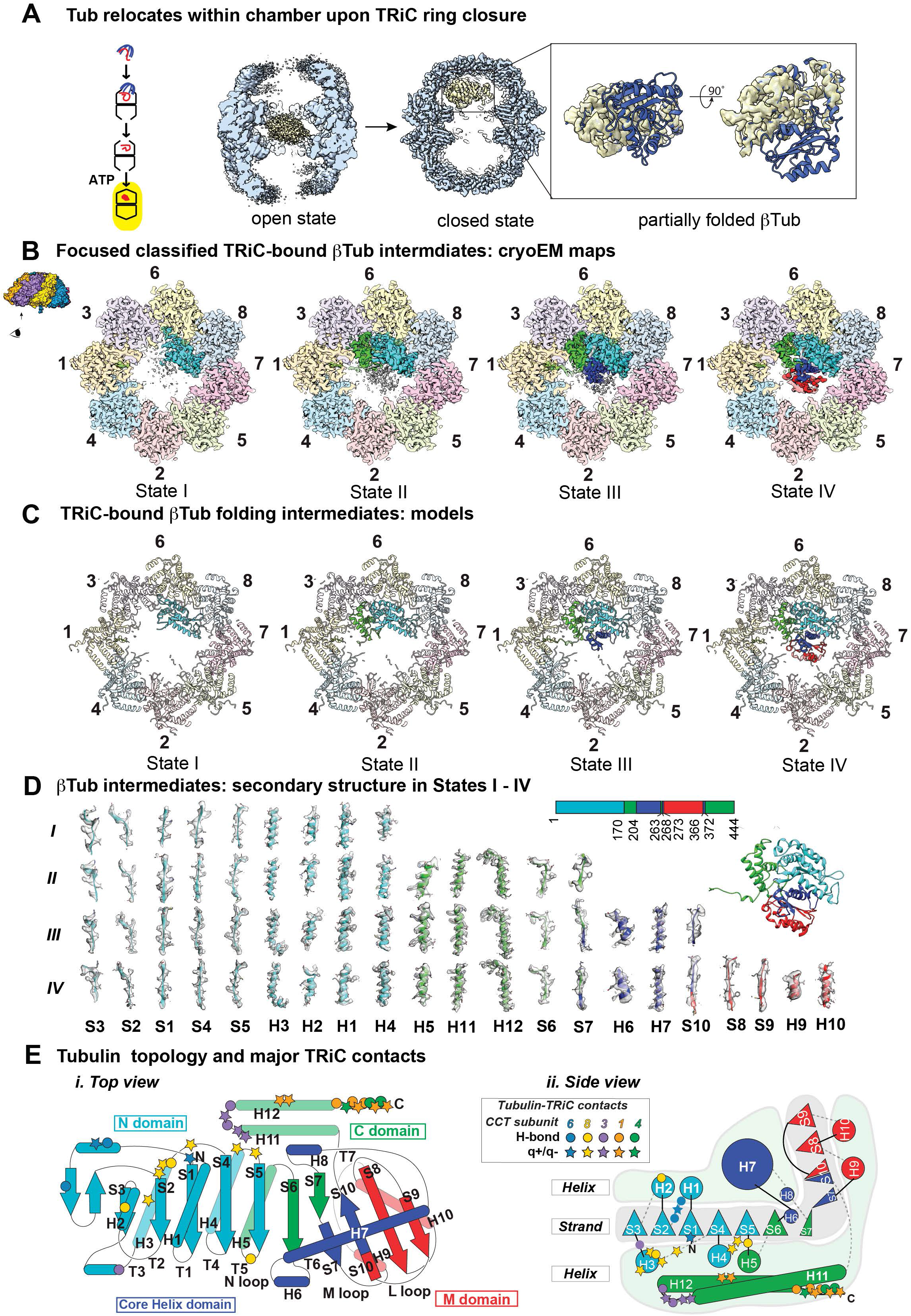
CryoEM identifies βTub folding intermediates in ATP-closed TRiC chamber. (A) CryoEM consensus maps of open and closed TRiC-βTub indicates dramatic repositioning of βTub density. (B) Focused classification on βTub identified four distinct states I-IV that could be separated by the amount of progressively folded βTub relative to the final state of βTub. (C) Atomic models of TRiC bound βTub folding intermediates.(D) Local resolvability of secondary structures in βTub folding intermediates reveals progressive formation of folded domains. (E) Cartoon illustration of βTub topology highlighting major contacts with TRiC subunits.

For each progressively formed βTub folding intermediate, we observed clearly defined density for secondary structure elements in the maps as well as clear density for the negatively charged C-terminal βTub tail, often called the E-hook (Figure 4D, S5A). We used a per-residue Q-score profile in a sliding window of 10 residues to examine the four identified TRiC-bound tubulin conformational states (Figure S4F) (Pintille et al, 2020). We obtained consistently continuous high Q-scores for the folded regions in each state (Pintille et al, 2020), demonstrating the high resolvability of structural elements in the folded domains of each state, in good agreement with their estimated resolution. In contrast, the poorly resolved segments have low Q-scores, suggestive of unfolded or conformationally dynamic regions. These conformational states, herein designated States I–IV, reveal progressively folded βTub conformations inside closed TRiC, whereby State IV corresponds to fully folded βTub (Movie S1). Notably, analysis of well-resolved residues in each βTub state map to specific domains formed by discontinuous sequences suggestive of domain-wise folding of tubulin inside the chamber (Figure 4D,4E). The least folded intermediate, herein State I, only contains the folded N-terminal βTub domain (residues 1-170) which is bound to the inner wall of CCT6-CCT8 (Fig. 4E). State I also has the C terminal tubulin E-Hook (residue 438-444) bound to a pocket between CCT1 and CCT4 (Fig. S5A). In addition to the folded N-terminal domain, State II contains the folded C-terminal domain (C-domain) of βTub, comprising resolvable discontinuous sequence elements, spanning residues 171-203; 263-267 and C-terminal helices 372-426. Of note, this folded domain was also bound to the inner wall of TRiC with residues of subunits CCT1 and CCT3 (Fig. 4E). State III contained, in addition to the folded N-and C-domains, the folded helical Core-domain, which also comprises discontinuous resolvable sequence elements spanning residues 204-262, 268-272 and 366-371. Finally, State IV additionally contains the folded middle M-domain, composed of residues 273-365, completing the fully folded βTub monomer. Though the TRiC-bound tubulin states exhibit C-alpha backbone variation in certain loop regions, the folded secondary structural elements in all four states are almost identical to the structure of native βTub (PDB 6I2I) (Figure S4G). Of note, the C-terminal E-hook tail of βTub was clearly traceable in all four states nestled in a pocket formed by CCT1 and CCT4 (Figure S5A). These four structural states of βTub within the chamber of TRiC suggest a folding pathway where discontinuous sequence elements in the encapsulated βTub polypeptide fold progressively in close association with the inner chaperonin chamber into specific domains to reach the native state (Figure 4E). We next examined in closer detail the nature of the βTub-TRiC chamber interactions.

### Domain specific CCT contacts spatially orient and restrain tubulin within the TRiC chamber

Upon ATP-driven encapsulation, we find that the inner TRiC wall engages in specific contacts with βTub intermediates. Of note, these interactions persist throughout the entire folding pathway. This suggests a very different mechanism than that observed for the bacterial chaperonin GroEL-ES, which promotes substrate folding by releasing encapsulated polypeptides into their central hydrophilic chamber (reviewed in Hartl et al., 2011), which has been often termed an “Anfinsen cage” (Ellis, 1996).

To understand how TRiC assists βTub folding, we examined the molecular contacts between its inner chamber and the encapsulated βTub. All four progressively folded βTub intermediates contain an already folded N-terminal domain, contacting specific residues in CCT6 and CCT8, as well as the very negatively charged, normally unstructured, C-terminal tail of βTub, also called the E-hook (Nogales et al., 1998b) nestled into a positively charged pocket at the interface of CCT1 and CCT4 (Figure 5B, S5A, Movie S2 and S3). States II-IV of βTub also contain the folded C-terminal domain of βTub contacting residues in CCT1 and CCT3. These contacts, which persist even when βTub reaches the fully folded state (Figure 5A), encompass an extended positively charged surface inside the TRiC chamber, formed by residues in CCT1-CCT3-CCT6-CCT8, and a complementary negatively charged continuous surface exposed upon folding of the N- and C-terminal domains of βTub (Figure 5B, Movie S3). Thus, electrostatic interactions with TRiC are progressively extended throughout the βTub folding pathway.

**Figure 5:**
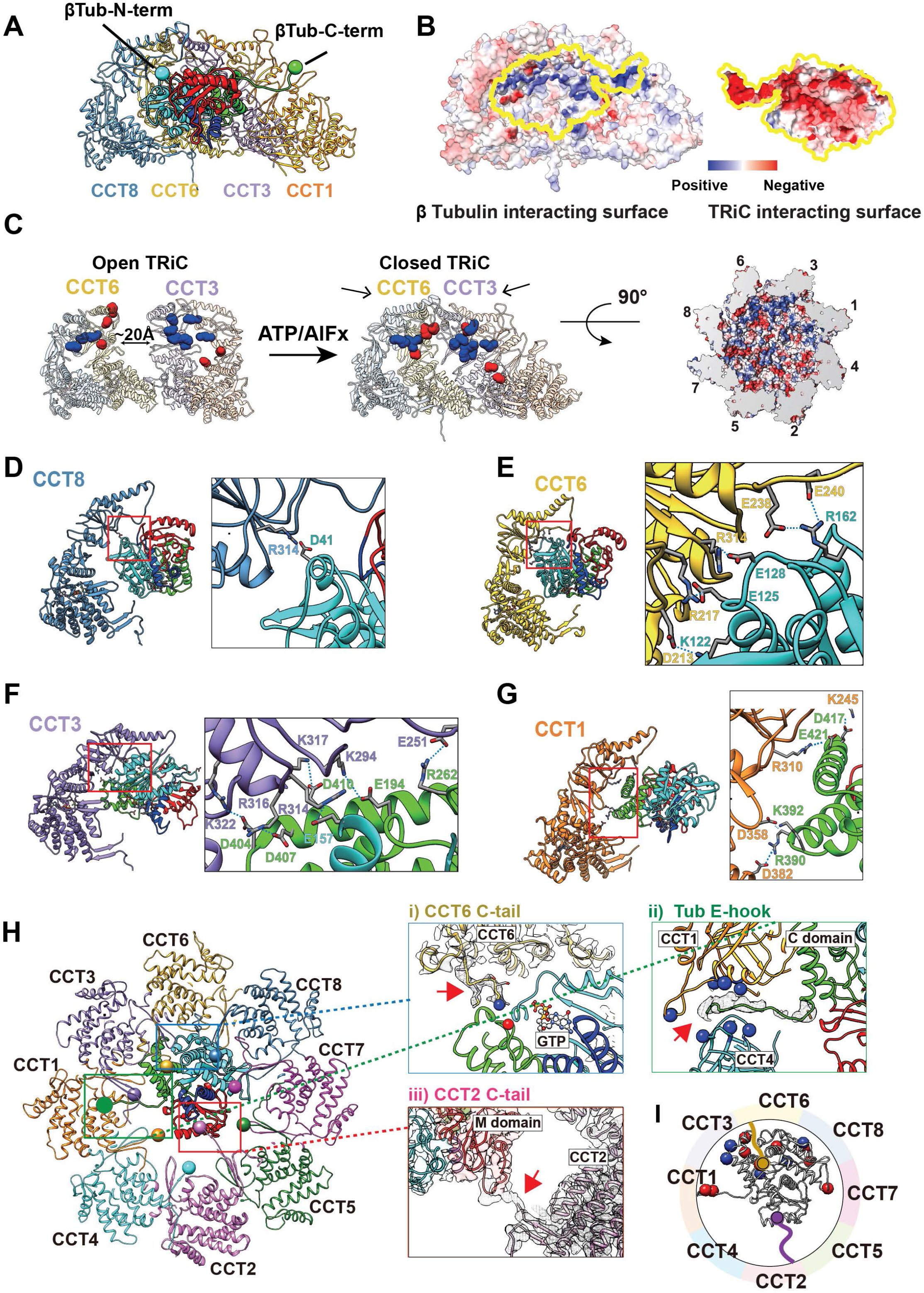
TRiC and βTub interaction in a domain specific manner. (A) Side view of a ribbon diagram of βTub and the CCT subunits (1,3,6,8) that have a large contact interface. The βTub N- and C-termini are indicated as a cyan and green ball, respectively. (B) The surface electrostatic distribution of the interacting surfaces of CCT (1,3,6,8) and βTub. (C) The formation of the electrostatic patch inner chamber of CCT (1,3,6,8) induced by the ring closure and the asymmetric charge distribution. (D-G) Side view of CCT (1,3,6,8) and βTub. βTub is colored by folding domains. Residues making salt bridges are labeled and sidechains are displayed. Each contact is shown as a dotted line. (H) Specific tail contacts between TRiC and βTub. Magnified views are displayed for (i) the CCT6 C-tail and βTub GTP binding pocket, (ii) βTub CCT1/4 binding pocket and βTub C-terminal E-hook tail, and (iii) CCT2 C-tail and βTub M-domain. (I) Simplified schematic of βTub, negatively charged residues (red) and positively charged residues (blue) that make contact with TRiC are indicated by spheres, the CCT2 (purple) and CCT6 (gold) C-terminal tails are also displayed with spheres on the terminal residue.

Previous analysis recognized that the TRiC inner chamber contains an asymmetric charge distribution (Movie S2), with a strong positively charged inner surface contributed by subunits CCT1/3/6/8, and a strong negatively charged inner surface contributed by CCT2/4/5/7 (Leitner et al., 2012) (Figure 5C). Of note, these charged surface patches are only formed upon lid closure following ATP hydrolysis (Figure 5C). The βTub electrostatic interactions with the positively charged surface of TRiC segregate folding to one side of the TRiC chamber, while the later folded regions of the polypeptide extend into the chamber cavity (Figure 4B, S5B). Notably, the density of these yet-to-be-folded dynamic segments appears to interact with the tail of CCT2 extending into the inner chamber space (Figure S5B), suggesting additional TRiC regions contribute to tubulin folding.

Our structural analyses of TRiC-βTub states reveal specific contacts between the CCT subunits and each βTub folding intermediates (Figure 5D -I). In the folded βTub N domain (blue in Figure 5D, E), residues on the N terminus, H2, H3, H4, and loops H1-S2, H3-S4 and H4-S5 form salt bridges and hydrogen bonds with residues on CCT6 and CCT8 (Figure 5D, E). Within the folded C-domain (green in Figure 5F,G), H11 and H12 are anchored to CCT1 and CCT3 by additional salt bridges and hydrogen bonds (Figure 4E). Our analysis also revealed specific contacts between βTub and tails of CCT6 and CCT2, in addition to the previously highlighted E-hook tail of βTub (Figure 5H, S5B, Movie S4). As both the E-hook and the CCT tails are normally disordered, it appears that formation of the TRiC-βTub complex promotes stabilization of these flexible regions. The CCT6 C-terminal tail is proximal to the βTub T5 loop in the C domain, which is part of the tubulin GTP binding pocket (Figure 5H-i). Since the T5 loop forms a possible hydrogen bond to GTP, and given that we already observe nucleotide density in State III of βTub, it is possible that the CCT6 tail facilitates formation of the tubulin GTP binding pocket. As discussed, the negatively charged E-hook of βTub is nestled into a positively charged pocket at the interface of CCT1 and CCT4 (Figure 5H-ii, S5A).

Unlike the N- and C-domains, the regions corresponding to the Core and M-domains, extend into the center of the TRiC chamber without any appreciable interactions with the inner wall in any of the intermediates. Closer examination reveals however that TRiC contacts the Core and M-domains via the C-terminal tail of CCT2 throughout the entire βTub folding pathway (Figure 5Hiii; S5B). In State I and II, the C-terminus of CCT2 interacts with unfolded Core/M domain density, while in States III and IV the CCT2 tail engages the N and M loops in the taxol binding site as the Core and M domains form sequentially during folding (Figure S5B) (Orr et al., 2003). This suggests the C-terminal tail of CCT2 interacts with and assists the folding of the Core and M-domains of tubulin within the lumen of the chamber. Interestingly, although the flexible CCT2 C-tail is poorly resolved in substrate-free closed TRiC chamber, we observe at a lower threshold an association between the vicinal C-terminal tails of CCT1 and CCT2 which extend together into the central chamber (Figure S5E). Consistent with this observation, the C-terminal tail of CCT1 is negatively charged and that of CCT2 is positively charged and possesses several prolines that may rigidify this flexible region. It is interesting to speculate that the C-termini of CCT1-CCT2 jointly act as a fulcrum that, upon lid closure, facilitates the repositioning of tubulin to the vicinity of the CCT1-CCT3-CCT6-CCT8 hemisphere.

Taken together, these structures indicate that each progressively formed βTub folding intermediate interacts with closed TRiC through a combination of more rigid complementary electrostatic interactions with the chamber wall and flexible interactions via βTub and CCTs tails (Figure 5I). Importantly, the negatively charged surface character of the N- and C-terminal domains is conserved across α-, β- and γ−tubulins (Figure S5F), suggesting a similar mode of TRiC-facilitated folding. Furthermore, the residues within the TRiC chamber making both electrostatic inner chamber and tail contacts with tubulin are highly conserved emphasizing the importance of these contacts in the tubulin folding process (Figure S5G). It thus appears that TRiC does not release the tubulin polypeptide into the closed chamber for folding. Instead, the TRiC chamber binds tubulin throughout the folding process. TRiC contacts directly restrain and orient the N-domain and C terminal tail of tubulin to one hemisphere in the chamber, thus limiting tubulin’s degree of freedom and preventing its free rotation in the chamber. We next examined how these interactions shape the intermediates leading to tubulin folding.

### TRiC interactions shaping the tubulin folding landscape

Our structural analyses suggest TRiC directs the βTub folding pathway through subunit specific contacts with the substrate (Figure 6A). In State I, two sets of contacts appear to promote the correct native βTub topology. Thus, electrostatic contacts between the positively charged surface on CCT6-CCT8 with the negatively charged surface of the already folded N-domain and those between the negatively charged C-terminal E hook tail while a positively charged pocket formed by CCT1 and CCT4 brings the distal C-terminal E-hook into proximity to the folded βTub N-domain. In turn, the positively charged surface on CCT1-CCT3 anchors the negatively charged surface of the βTub C-domain, likely promoting its subsequent folding in State II and leading to formation of the N- and C-domain interface, in a structure highly homologous to the fully folded domains in native βTub. The molecular contacts with the TRiC inner wall are almost completed at this stage. The Core and M-domains do not exhibit any interactions with the wall of TRiC but rather face the interior of the chamber. However, the C-terminus of CCT2 engages the Core and M-domains throughout the folding process, starting with the unstructured polypeptide in States I and II, and continuing as these domains fold sequentially in States III and IV. We speculate the intrinsically disordered CCT2 tail may directly assist the folding of these two domains, contributing to TRiC chaperone function during the folding reaction. The early folding of the N-terminus in TRiC-bound βTub resonates with the PK-sensitivity assays of the folding reaction, which indicates early protection of an N-terminal domain, but rapid digestion of the C-terminus until completion of full-length βTub folding (Figure S1G-I).

**Figure 6:**
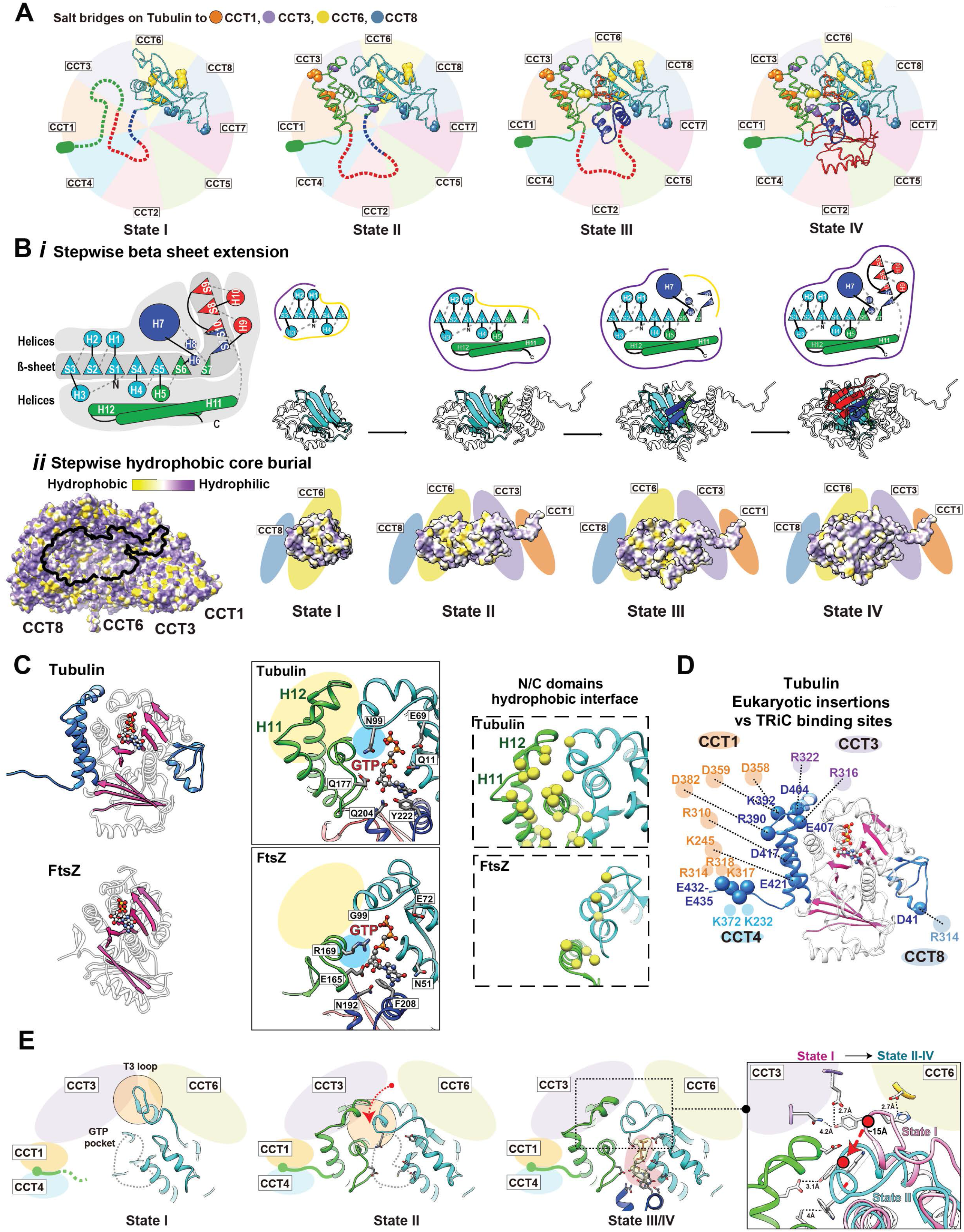
The directed folding of βTub intermediates by specific CCT interactions. (A) Cartoon showing specific contacts between βTub domains and CCT subunits at each intermediate state. βTub residues that make salt bridges are shown as balls and colored according to the interacting CCT subunit. The missing domains of each state are shown as dotted lines. The negatively charged C terminus is represented as a green ellipse. (B)(I) Cartoon of βTub to emphasize the hydrophobic β-sheet running the length of the βTub structure that is surrounded by α−helices, and individual cartoons for the sequential formation of this β-sheet core through each folding state. (ii) Hydrophobicity distribution on TRiC of the βTub interacting surface, and of the hydrophobic core burial of βTub as it extends towards the interior of the chamber through the folding states. (C) Ribbon diagram of βTub (PDB 6I2I, top panel) and FtsZ (PDB 1FSZ; bottom panel) with additions unique to βTub structure shown in blue. The overall β-strand folds in βTub and FtsZ are colored in pink. Zoom-in views of the GTP binding pocket (contacting residues indicated) and N/C-domain interface (hydrophobic residues yellow) are shown for comparison. (D) The contacts between βTub specific insertions and CCT subunits are shown with potential salt bridges. (E) Formation of the βTub GTP binding pocket through βTub intermediates displayed in ribbon diagrams. The GTP pocket is indicated as a gray dotted line in states I and II, and the density map corresponding to the nucleotide is displayed as a yellow density with pink background in state III/IV. The highlighted T3 loop makes a large conformational change between states I and the others indicated by the red arrow and displayed in the focused overlay between states I and II-IV.

Our analysis suggests TRiC directs tubulin folding to circumvent challenges posed by its complex topology and aggregation-prone β-sheets. Thus, while TRiC-directed domain-wise folding is discontinuous in the βTub sequence, it orchestrates the sequential formation of a hydrophobic β-sheet running the length of the βTub structure (Figure 6Bi, Movie S5). Notably, anchoring the surface of the folded N- and C-domains to the TRiC chamber wall not only establishes the native topology but also positions the growing folding intermediates so that their hydrophobic core extends towards the interior of the chamber (Figure 6Bii). Each intermediate in the folding pathway adds another β-strand layer to the exposed hydrophobic core, priming it for the next folding step. Importantly, each step also incorporates the helices above and below the growing β-sheet thus protecting its edges and reducing the potential for off-pathway interactions. In State I, the folded N-domain exposes the hydrophobic strands S4 and S5. Completion of the Rossman fold in State II allows helices H5, H11 and H12 to cover the newly incorporated sheets S6 and S7 (Rossmann et al., 1974). Formation of the Core domain in State III requires a change in direction between strands S7 and S10, mediated by helix H7. The CCT2 tail may participate in this step, as it interacts with this region of the Core domain. The last folding step to reach native βTub, involved completion of the β-sheet by addition of sheets S8-S10 and positioning of helices H7, H9 and H10 to cover the hydrophobic core of tubulin. Accordingly, the TRiC interactions shape the tubulin folding landscape to preclude the formation of trapped folding intermediates.

### TRiC action in folding pathway in the context of tubulin evolution

While tubulin folding exhibits an obligate requirement for TRiC, its prokaryotic homolog FtsZ can spontaneously fold without chaperones (Andreu et al., 2002; Erickson et al., 2010). Both proteins share a similar overall topology with other superfamily members in prokaryotes and archaea, including the overall domain organization and the extended core beta sheet (Figure 6C; S6A-B) (Andreu et al., 2002; Aylett and Duggin, 2017; Nogales et al., 1998a; Yutin and Koonin, 2012). However, eukaryotic tubulins are distinguished by additional features integral to their unique function and regulation, including remodeling of GTP affinity and hydrolysis kinetics; ability to engage in microtubule lateral contacts; regulation by post translational modifications (PTMs) and binding to kinesin and microtubule associated proteins (MAPs) (Andreu et al., 2002; Janke and Bulinski, 2011; Nogales et al., 1998b; Skiniotis et al., 2004). These tubulin functions are accompanied by insertions in the H1-S2 loop and two negatively charged helices, H11 and H12; by the presence of the E-hook tail; by the remodeling of the residues contacting GTP, and by extensive changes to the C-domain leading to an increase in hydrophobic interdomain interfaces (Andreu et al., 2002; Aylett and Duggin, 2017; Nogales et al., 1998a; Yutin and Koonin, 2012) (highlighted in Figure 6C for 6I2I vs 1FSZ, S6C). Eukaryotic tubulins are also distinguished by the strongly negative surface charge on the folded N- and C-domains as well as a negatively charged C-terminal E-hook tail. We hypothesize that these features of eukaryotic tubulins compromise their ability to fold either spontaneously or even with simpler chaperone systems. Remarkably, the tubulin-specific insertions correspond to the sites of TRiC binding during folding in the closed chamber, suggesting their incorporation into an ancestral tubulin was accompanied by a need for specialized folding assistance (Figure 6D).

We delved into this hypothesis by close examination of how TRiC mediates formation of the unique βTub N-C-domain interface (Figure 6E, S6D). The T3 loop that is part of the N-C interdomain interface in folded tubulin shifts between two alternative conformations as TRiC-bound tubulin folding progresses, with 13 Å C-alpha variation (Fig. S4G, S5C, Movie S6). Our maps resolve the location of bulky side chains in the T3 loop, such as H105 and Y106, allowing us to trace the density of T3 along the folding pathway with valid Q scores (Figure S6D). The H1-S2 loop forms charged and hydrogen bonded interactions with the TRiC chamber suggesting a stabilizing effect on the inserted tubulin loops by the TRiC chamber (Figure 4E, 6D). In State I, the T3 loop, resolved at residues 100-109 adopts an extended conformation directed by contacts with CCT3 and CCT6 (Fig. S5C). In particular, βTub H105 is proximal to D86 of CCT6, and βTub Y106 is possibly hydrogen bonded with E202 of CCT3 (Figure 6E). In State II, a decrease in resolvability of residues 96-106 in T3 (Fig. S5C), suggests the loop changes conformation upon folding of the C-domain. In States III and IV, the T3 loop becomes well resolved due to its stabilization by interaction with the folded C-domain, adopting the native βTub conformation (PDB: 6I2I) (Figure S4G, S5C). Accordingly, N99 of T3 serves in GTP γ-phosphate sensing while hydrophobic T3 residues including W101 are embedded in the hydrophobic interface between the N and C domains (Figure 6C). Folding of the core helix domain completes the GTP binding pocket by contributing the H6-H7, consistent with the nucleotide density in the GTP binding pocket observed in States III and IV (Fig. S5C,5D, Figure 6E). These observations indicate that TRiC contacts guide the T3 loop from an extended conformation before C-domain folding to the native conformation after the C domain is folded.

Our analyses are consistent with previous studies indicating GTP binding contributes to tubulin stability (Zabala et al., 1996) and previous genetic analyses of tubulin mutants (Figure S6E) showing mutations in the N/C-domain interface affect βTub folding and release from TRiC (Wang et al., 2006). We conclude that TRiC contacts directly guide the proper progression of events needed to form the GTP binding pocket of βTub, by sequential positioning of the nucleotide sensing T3 loop to form the N-C domain interface, which are not conserved in FtsZ (Figure 6C-E1). Collectively, TRiC may thus have coevolved to support the function and regulation of βTub by actively assisting and directing its folding pathway.

## DISCUSSION

### The PFD-TRiC pathway steers βTub folding through specific intermediates

Newly translated tubulin folds assisted by chaperones PFD and TRiC/CCT (Figure 7A) (Geissler et al., 1998; Vainberg et al., 1998). How TRiC, promotes folding has long been enigmatic, as proteins such as tubulin have an obligate requirement for TRiC assistance to fold. Overviewing the tubulin folding mechanism revealed by biochemical and structural experiments in this study, we find PFD maintains unfolded states in the bound substrate and directly delivers it to TRiC. This promotes compaction of the substrate, which engages in extensive interactions with the CCT tails. Upon ATP hydrolysis, specific interactions with individual CCT regions inside the closed TRiC chamber guide tubulin to the native state through stepwise formation of partially folded intermediates (Figure 7B). Throughout this pathway the substrate is not released into the central TRiC chamber, unlike what is observed with GroEL-ES, but instead remains anchored through specific contacts with the CCT wall and the C-tails. These findings establish that, for tubulin, and perhaps other TRiC substrates, the folding pathway is guided by explicit interactions with the inner TRiC chamber.

**Figure 7:**
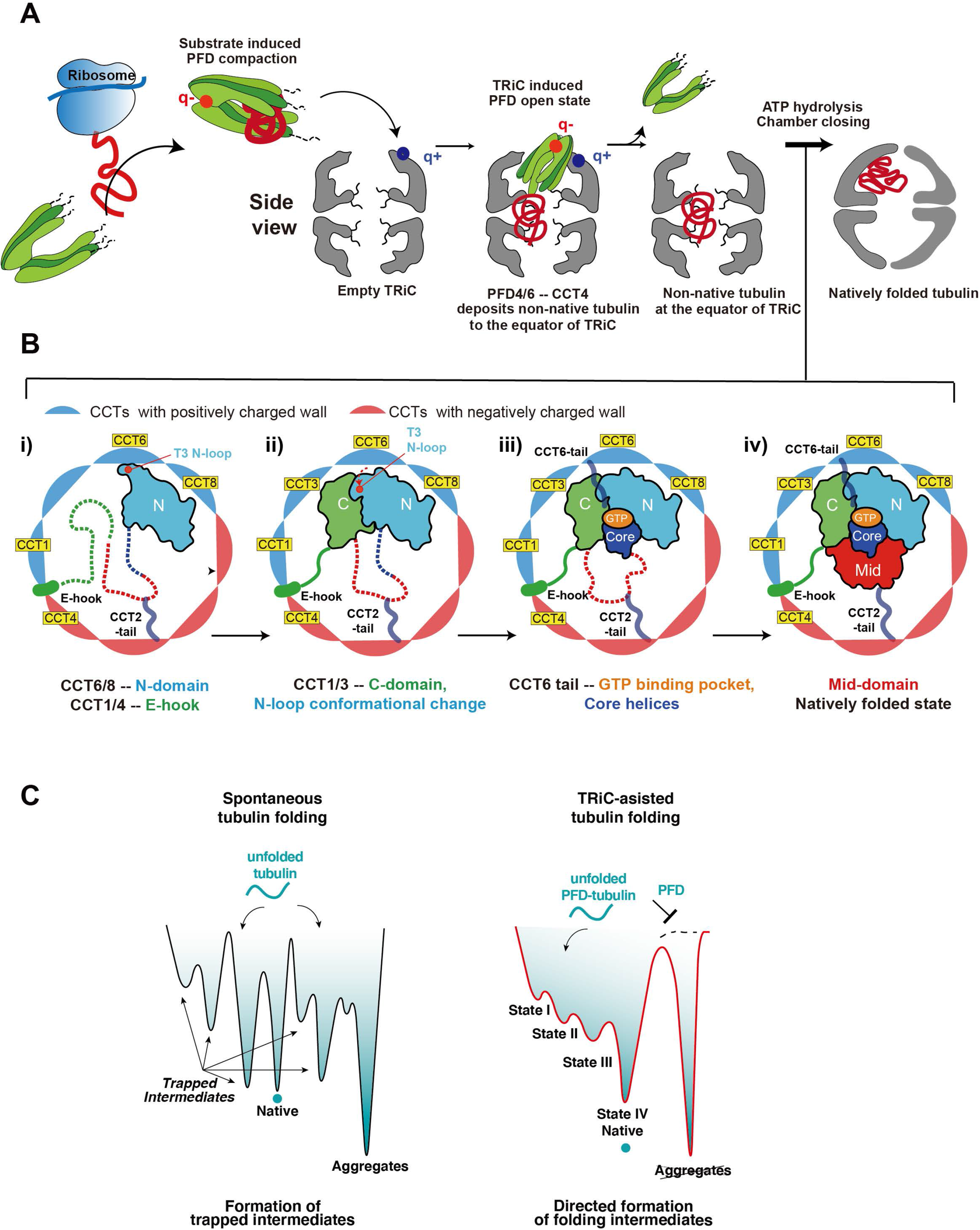
Mechanism of TRiC directed folding pathway for βTub. (A) Model of βTub folding pathway; Nascent chain βTub translated from the ribosome is captured by PFD. The binding induces the structural compaction of PFD and βTub remains in an unfolded, soluble state. Red and blue dots on PFD and TRiC show key electrostatic points mediating initial PFD binding and repositioning into the chamber. TRiC captures βTub from PFD in the third chamber illustrated with the unresolved tails in the third chamber with PFD transitioning to an open, low-substrate affinity, conformation. βTub capture by TRiC induces a partially folded βTub at the TRiC equator. ATP hydrolysis induces closure of TRiC and the substrate repositions into the chamber while undergoing folding. (B) Upon chamber closure, TRiC directs the folding pathway of βTub through the distinct intermediates; unfolded domains are shown in dotted line and CCT subunits are colored according to their charged state. (i) The N-domain is anchored to the positively charged surface of CCT6/8. The E hook is held by the positively charged pocket from CCT1/4 and the T3 loop is stabilized in the extended conformation through interactions with CCT3/6. (ii) The βTub C-domain folds and is stabilized by CCT1/3. The T3 loop repositions off the TRiC chamber wall forming a stable interface between the βTub N- and C- domain, which promotes GTP binding pocket formation. (iii) The Core domain completes the formation of the GTP binding pocket and the nucleotide binding provides additional stabilization. (iv) The M domain folds last completing the native tubulin fold. Note that the tails of CCT2 and CCT6 engage throughout the folding process by direct contacts to the GTP binding pocket and the M domain of β-tub, respectively. (C) Schematic folding landscape of tubulin folding spontaneously (left) or directed by TRiC (right). Spontaneous tubulin folding leads to trapped intermediates and/or in aggregates often observed in folding assays. For TRiC/PFD assisted tubulin folding, PFD maintains tubulin unfolded and prevents aggregation. TRiC promotes folding by directing the formation of progressively folded intermediates and preventing off-pathway trapped intermediates.

### Defining PFD-βTub interactions and subsequent transfer to TRiC

βTub forms a complex with PFD during biogenesis (Hansen et al., 1999; Vainberg et al., 1998). We formed this complex by co-expression *in vivo* and found PFD maintains tubulin in an unstructured, highly protease sensitive conformation (Figure 1, S1). XL-MS analyses also indicate a highly dynamic interaction between tubulin and the inner surface of the coiled-coil delimited PFD chamber. Native mass spec and native gel analyses reveal that in the absence of substrate PFD alternates between compact and extended conformations. Binding of tubulin shifts PFD to the compact state. Previous structural analyses of the PFD conformers in PFD-TRiC complexes (Gestaut et al., 2019), together with our current analyses of PFD-TRiC-βTub, provide intriguing clues that these states are relevant to the mechanism of substrate handover from PFD to TRiC (Figure 2I-J, 2SE-F), These analyses suggest that PFD initially contacts TRiC in the compact, substrate-bound, state through contacts between the PFD4-PFD6 coils and the CCT4 apical domain (Gestaut et al., 2019). When the PFD and TRiC chambers become aligned, the PFD2 tentacles are repositioned, leading to the open conformation, facilitating substrate release. Our analysis of the TRiC-PFD-βTub ternary complex obtained here supports this hypothesis.

While PFD-βTub is stable in solution, TRiC addition causes a rapid transfer of βTub to TRiC. Consistent with this, we detect extensive PFD-βTub crosslinks in the binary complex, but the ternary TRiC-PFD-βTub complex yield almost exclusively βTub crosslinks to TRiC, indicative of a transfer of substrate to the chaperonin chamber. Visualization of TRiC-PFD-βTub shows PFD in the open state and substrate density in the TRiC chamber. We hypothesize substrate transfer from PFD to TRiC happens in two steps; first PFD recognizes TRiC through an electrostatic interaction, and second PFD coiled-coils pivot to align the TRiC/PFD chambers. In this fully engaged complex with TRiC, PFD adopts the open, low substrate affinity state allowing substrate transfer from PFD to TRiC (Figure 7A).

### The disordered chaperonin tails enmesh βTubulin within the open TRiC chamber

Upon PFD delivery to TRiC, βTub becomes more protected from protease digestion suggesting a more compact state. The substrate appears to become compacted within the TRiC chamber via extensive interactions with the intrinsically disordered N- and C-termini of TRiC. Indeed, XL-MS analysis identified significant contacts happen through the intrinsically disordered terminal tails of PFD and TRiC with the early βTub folding intermediate. Surprisingly, most of the density attributable to tubulin is in a central inter-ring chamber that appears to also contain the intrinsically disordered N- and C-terminal CCT tails. We did not observe substrate density in the apical domains of TRiC, unlike a recent study of TRiC-actin in the open state (Balchin et al., 2018). It is possible that the interactions of βTub with the apical domains are too dynamic to be resolved at high resolution. Alternatively, βTub could be initially captured by the apical domains and then transferred to the inner chamber by the interaction with the CCT tails. Supporting this idea, a previous study observed crosslinks between βTub and specific sites in the apical domains of TRiC (Joachimiak et al., 2014).

Analysis of βTub directly bound to TRiC also reveals substrate density in the central chamber similar to what is observed upon transfer from PFD. No clear folded domain or even secondary structure is found in the cryoEM map density attributable to βTub. Of note, no density in the central inter-ring chamber is observed without βTub addition, suggesting interaction of the substrate with the CCT tails generates a dense compact entity. Possibly the CCT tails are not observed in any high resolution TRiC/CCT structure due to their flexibility, even though they collectively constitute approximately 35.5 kDa of protein. Our findings contrast a previous study in which a ∼4.0 Å map of TRiC bound to mLST8 (Cuellar 2019) was interpreted as having clear secondary structure similar to the final β-propeller folded protein; however, the density was not well resolved and the folded state of the substrate was not confirmed in any biochemical assay. The exact conformation of the substrate within this TRiC chamber remains to be determined. One intriguing possibility is that the disordered tails confined within the chamber act as a tethered solvent to maintain the fluidity of the substrate promoting a dynamic yet compacted state. The non-native polypeptide would engage in polyvalent low affinity interactions with the distinct polar and non-polar motifs in the different CCT tails. The liquid like nature of such a coacervate may prevent the formation of non-productive or trapped intermediates, priming the substrate for productive folding upon chamber closure.

### Beyond Anfinsen: the TRiC chamber shapes folding intermediates

Upon TRIC ATP hydrolysis, PFD leaves and the TRiC chamber is closed. Lid closure repositions the substrate from the equatorial chamber into one of the ring-enclosed folding chambers (Figure 7B). CryoEM provided unprecedented insight into the folding process of βTub inside the ATP-closed TRiC chamber. Perhaps most surprising is the visualization of stepwise βTub folding through intermediates that remain bound to specific regions in the chamber. Initially, βTub makes contact with the TRiC chamber through electrostatic and H-bond interactions from the βTub N-domain to CCT6 and CCT8, and the negatively charged C-terminal βTub E-hook contacting a distal positively charged pocket between CCT1 and 4 (State I). These contacts establish the necessary native topology for βTub, allowing folding of the C-domain and subsequent formation of the N-C-domain interface (State II). Both N- and C-folded domains remain anchored to the TRiC wall throughout the folding process. We hypothesize that the highly hydrophilic surface of the TRiC chamber together with interaction with polar CCT tails helps orchestrate a hydrophobic collapse to then drive sequential folding of the Core and Middle domains of βTub (State III and IV). This TRiC-mediated folding enables formation of the GTP binding pocket and binding to nucleotide in State III. One feature of the discontinuous formation of tubulin domains in this folding pathway is the unidirectional assembly of the β-sheet spanning the entire hydrophobic core in folded βTub. These TRiC contacts appear to direct βTub folding through sequential domain formation without releasing the substrate into the chamber. This is in contrast to what is observed for folding by the chaperonin, GroEL,-ES which releases polypeptides into an inner chamber which provides a unique confined environment for folding (Chen et al., 2013; Fenton and Horwich, 1997; Gupta et al., 2014; Motojima, 2015; Sharma et al., 2008).

Our analysis strongly suggests that TRiC uniquely and specifically alters the βTub folding pathway (Figure 7C). It is clear that, at least for tubulin, the closed chamber is not an “Anfinsen cage”, unlike what is observed for bacterial chaperonins, which do release the substrate for folding within the ATP-closed chamber (Sharma et al., 2008) Since tubulin does not refold reversibly even at low concentrations or temperatures (Andreu et al., 2002) it is not clear whether reaching the native state is under thermodynamic control. Interestingly, biochemical studies (Andreu et al., 2002) as well as molecular dynamics simulations of unfolding (Dima and Joshi, 2008) have shown that the N and C-terminal domain interface in tubulin is the least stable and the first to unfold, leading to loss of nucleotide binding, while the M and Core domains are the most stable and last to unfold (Dima and Joshi, 2008). Notably, TRiC guided folding upends the expectation that these more stable domains form first. Instead, the TRiC chamber prevents initial formation of the more stable M- and Core domains and prioritizes establishment of the correct topology by simultaneous electrostatic binding of the C-terminal E-hook tail and folded N domain, followed by folding of the C-domain. The fact that contacts with the closed TRiC chamber guide multiple steps in βTub folding raises questions on the universality of Anfinsen’s Principle.

The concept of TRiC directing the folding pathway is illustrated by the notable structural change in the T3-loop of βTub between States I and II. In State I, TRiC keeps the T3-loop in an extended conformation through contacts with the inner wall (Figure 7B). In State II, the C-terminal domain folds through association with the TRiC wall, while the T3-loop is released from the TRiC wall to become nestled at the N-C domain interface. The T3 loop then acts as a folding switch to control sequential folding of the GTP binding pocket after completion of the N-C domain interface (Figure 6E). Thus, TRiC binding to the T3-loop ensures the proper folding of the C-domain, establishing the overall fold topology of βTub and allowing the hydrophobic core of the M and Core domains to fold. Perhaps GTP binding to the incipient nucleotide pocket drives these subsequent steps, similar to what is observed for bacterial Tubulin FtsZ (Figure 6C), where GTP binding is a folding limiting step required for spontaneous refolding (Huecas et al., 2020). While our analyses strongly suggests that TRiC directs the formation of specific folding intermediates (Figure 7B), it is also possible that the contacts of the substrate with the chamber serve to disfavor formation of kinetically trapped states (Figure 7C).

### Insights into evolution of eukaryotic proteins that cannot fold without TRiC

It is intriguing to speculate why and how this complex chaperonin requirement evolved from simpler ancestral proteins like FtsZ, which can refold without assistance. Compared with FtsZ, tubulin contains a series of structural changes which serve functionally to enable homotypic interactions at the lateral interface of protofilaments and binding of microtubules associated proteins (MAPs) and Kinesin (Bertrand et al., 2005; Nogales et al., 1998b; Skiniotis et al., 2004). While these additional elements support the novel properties of eukaryotic tubulin, they appear to impair spontaneous folding (Bertrand et al., 2005), perhaps leading to off-pathway or kinetically trapped intermediates (Figure 7C). Strikingly, the major sites of βTub interaction with the TRiC-chamber are in these eukaryote-specific regions. We propose these eukaryotic-specific changes in tubulin evolved to exploit the hetero-oligomeric and asymmetric nature of TRiC to circumvent the inherent challenges they pose to the folding pathway.

Our study raises new questions about tubulin biogenesis. For βTub, but also likely for αTub, downstream tubulin dimer assembly factors are not required to reach the native folded state. It will be of interest to determine how folded TRiC-bound tubulin monomers are released from the chaperonin. Perhaps downstream factors involved in α, βTub dimer formation interact with TRiC to disrupt the tubulin binding sites, alternatively TRiC reopening during ATPase cycling break up the tubulin-binding sites in the chamber allowing for passive binding of the cofactors. The finding that tubulin folding is closely directed by TRiC chamber interactions opens new perspectives to understand the role of TRiC in eukaryotic protein evolution. The striking expansion of TRiC subunits directs the subunit-specific diversification of substrate binding sites, ATP affinity and chamber properties all of which contribute to its unique ability to fold some eukaryotic proteins. In turn, these may underlie their obligate requirement for TRiC to fold. Given that TRiC facilitates folding of ∼10% of the proteome, it is likely that some proteins employ TRiC like an Anfinsen Cage, similar to that of GroEL-ES, that prevents aggregation and smooths the folding landscape (Georgescauld et al., 2014). Other proteins may exploit other evolved features of the TRiC hetero-oligomer. For instance, the diverse apical domain substrate specificity may promote a specific polypeptide binding topology (Balchin et al., 2018; Joachimiak et al., 2014; Roh et al., 2016; Spiess et al., 2006). Also, some substrates may interact distinctly with other regions in the chamber (Knowlton et al., 2021). Our finding that the eukaryote-specific tubulin insertions evolved surface-exposed motifs that bind specific regions in the TRiC chamber to shape the folding pathway provides an intriguing clue as to how some eukaryotic proteins evolved to exploit the unique properties of TRiC to acquire new functionalities that would otherwise impede their correct folding.

## ACKNOWLEDGMENTS

This work has been supported by the National Institutes of Health grants R01GM074074 to JF; P41GM103832, S10OD021600 and R01GM079429 to WC and the Korean National Research Foundation (2019R1C1C1004598, 2020R1A5A1018081, 2021M3A9I4021220) and SUHF foundation to SHR. The cryo-EM data was collected at the Stanford-SLAC CryoEM facilities. We thank members of the Frydman lab for useful discussions and suggestions. We thank Dr. P. Picotti and the UCSF Mass Spectrometry Facility for access to instrumentation.

## AUTHOR CONTRIBUTIONS

JF, DG, WC and SHR conceived the project; DG carried out all cloning, protein purifications and biochemical experiments; BM collected cryo-EM datasets of PFD-TRiC-Tubulin and TRiC-Tubulin with ATP/AlFx and performed initial data analysis. YZ collected the apo TRiC data and carried out cryo-EM data analysis and focused classification of the TRiC-Tubulin under ATP/AlFx condition; JP carried out cryo-EM data analysis on PFD-TRiC-Tubulin in the open state and apo TRiC state. DG and AL carried out the XL-MS experiments and analyzed data for all complexes; MC carried out native mass spectrometry; GP generated Movie S1. DG, YZ, J.F, JP, WC, SHR wrote the MS. All authors contributed to the final MS.

