## Supplementary info for "Structural visualization of the tubulin folding pathway directed by eukaryotic chaperonin TRiC"

#### LIST OF SUPPLEMENTAL INFORMATION

Figure S1. Purification systems and folding analysis for α/β Tubulin; Related to figure 1
Figure S2. Native mass and XL analysis for PFD and PFD-βTub; Related to figure 2
Figure S3. CryoEM workflow and XL analysis for PFD-βTub-TRiC; Related to figure 3
Figure S4. CryoEM workflow and analysis for TRiC-βTub; Related to figure 4
Figure S5. TRiC chamber spatially orient and restrain βTub; Related to figure 5
Figure S6. Tubulin coevolved with TRiC for directed folding process; Related to figure 6.

Movie S1. Cryo-EM density of the four βTub folding states in closed TRiC chamber; Related to figure 4
Movie S2. Asymmetric charged surface of TRiC closed chamber; Related to figure 5
Movie S3. Electrostatic interaction between βTub N-, C-domain and the TRiC chamber wall; Related to figure 5
Movie S4. Description of the E hook tail of βTub and CCT2/6 C-terminal tail in the βTub folding process; Related to figure 5
Movie S5. Inter-domain hydrophobic burial in the βTub folding process and continuous hydrophobic framing of the growing β-sheet in the folding process; Related to figure 6
Movie S6. Conformational change of T3 loop from State I to the other states; Related to figure 6

Table S1. CryoEM data collection, 3-D reconstruction, and refinement - PFD-βTub-TRiC
Table S2. Crosslinking and mass spectrometry Data

### Figure S1

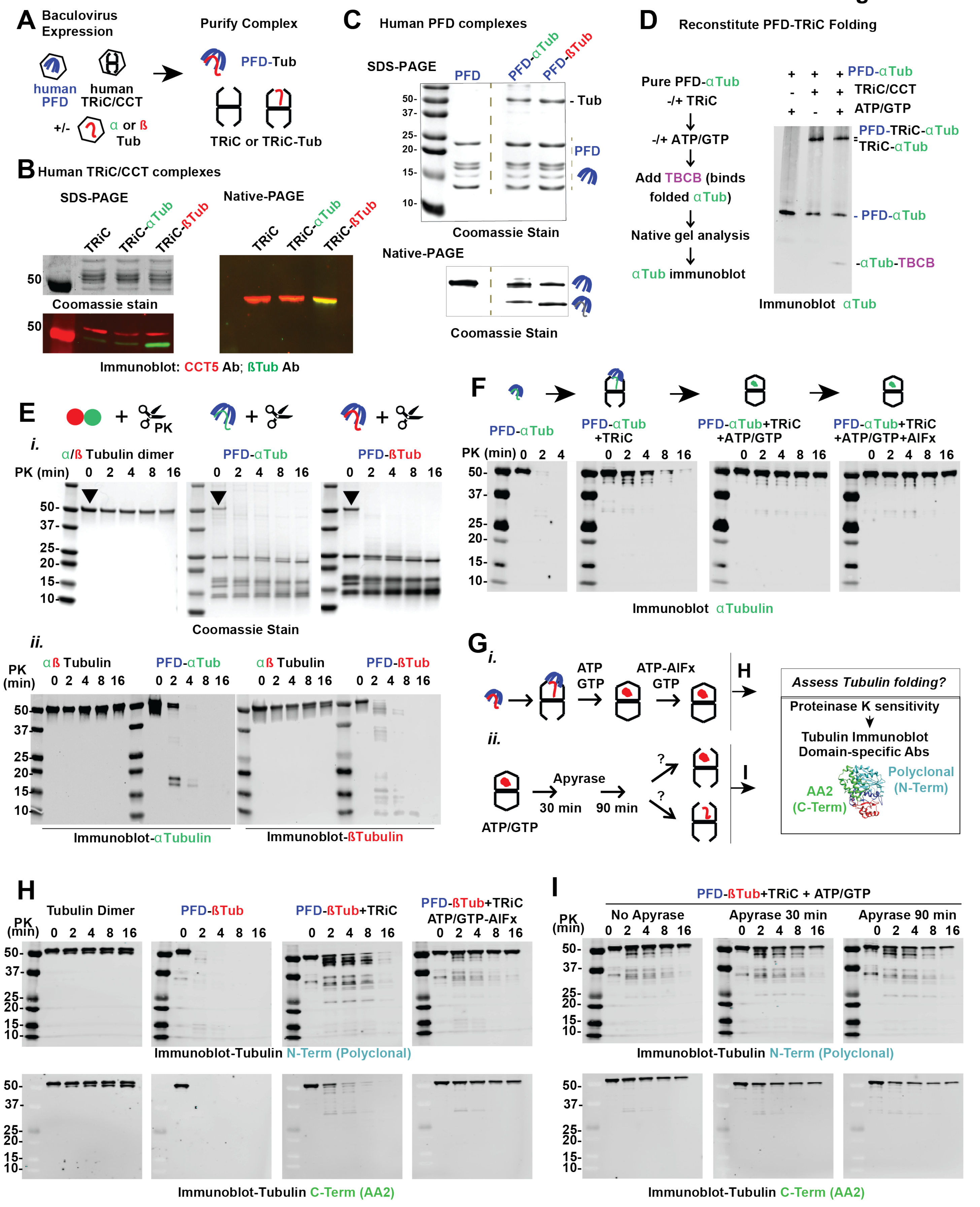

A Baculovirus Expression
human PFD
human TRiC/CCT
+/- α or ß Tub
Purify Complex
PFD-Tub
TRiC or TRiC-Tub
B Human TRiC/CCT complexes
SDS-PAGE
TRiC
TRiC-αTub
TRiC-ßTub
Coomassie stain
Native-PAGE
Immunoblot: CCT5 Ab; ßTub Ab
C Human PFD complexes
SDS-PAGE
PFD
PFD-αTub
PFD-ßTub
- Tub
PFD
Coomassie Stain
Native-PAGE
Coomassie Stain
D Reconstitute PFD-TRiC Folding
Pure PFD-αTub
-/+ TRiC
-/+ ATP/GTP
Add TBCB (binds folded αTub)
Native gel analysis
αTub immunoblot
PFD-αTub
TRiC/CCT
ATP/GTP
PFD-TRiC-αTub
TRiC-αTub
- PFD-αTub
-αTub-TBCB
Immunoblot αTub
E
PK
i.
α/ß Tubulin dimer
PFD-αTub
PFD-ßTub
PK (min) 0 2 4 8 16
Coomassie Stain
ii.
PK (min)
αß Tubulin
PFD-αTub
PFD-ßTub
Immunoblot-αTubulin
Immunoblot-ßTubulin
F
PFD-αTub
PFD-αTub +TRiC
PFD-αTub+TRiC +ATP/GTP
PFD-αTub+TRiC +ATP/GTP+AlFx
PK (min)
Immunoblot αTubulin
G
i.
ATP GTP
ATP-AlFx GTP
ii.
ATP/GTP
Apyrase
30 min
90 min
H
I
Assess Tubulin folding?
Proteinase K sensitivity
Tubulin Immunoblot Domain-specific Abs
Polyclonal (N-Term)
AA2 (C-Term)
H
Tubulin Dimer
PFD-ßTub
PFD-ßTub+TRiC
PFD-ßTub+TRiC ATP/GTP-AlFx
PK (min) 0 2 4 8 16
Immunoblot-Tubulin N-Term (Polyclonal)
Immunoblot-Tubulin C-Term (AA2)
I
PFD-ßTub+TRiC + ATP/GTP
No Apyrase
Apyrase 30 min
Apyrase 90 min
Immunoblot-Tubulin N-Term (Polyclonal)
Immunoblot-Tubulin C-Term (AA2)

**Figure S1. Purification systems and folding analysis for α/β Tubulin; Related to figure 1.**

(A) Purification scheme for PFD-Tub and TRiC-βTub (B) Purified TRiC-βTub or αTub with SDS-PAGE and Native-PAGE analysis, immunoblot confirms βTub or αTub co-migrates with TRiC in native gel. (C) Purified PFD-Tub (β or αTub) with SDS-PAGE and Native-PAGE analysis, note faster migration of PFD-Tub relative to PFD alone in native gel. (D) αTub folding assay. αTub co-expressed/purified with PFD comigrates on a native gel. In the absence of nucleotide αTub comigrates with TRiC, and addition of nucleotide (ATP/GTP) drives folding and partial release of αTub in the presence of TBCB. (E) Protease sensitivity assay (Proteinase K) of PFD-Tub samples; Fully folded α/β Tubulin dimer is insensitive to PK digestion, while PFD-Tub samples show rapid degradation by SDS-PAGE visualized by Coomassie stain (i) or immunoblot (ii). (F) Protease sensitivity assay of PFD-αTub folding by TRiC. Like βTub, αTub bound to PFD is protease sensitive, transfer to TRiC from PFD causes a decrease in sensitivity, and addition of nucleotide drives folding resulting in limited protease sensitivity equivalent to αTub in a native α/β Tub dimer. (G-I) Protease sensitivity assay to test if TRiC closure is providing protease protection with degradation assayed by both an N-terminal region polyclonal antibody, and a C-terminal (AA2) monoclonal antibody. After completion of βTub folding by TRiC as assayed in (H), apyrase is added to open the TRiC complex and protease sensitivity is assayed at indicated time points (I). No increase in protease sensitivity is observed immediately after apyrase addition, while 90 min after addition a mild increase in sensitivity is observed likely due to slow unfolding.

**Figure S2**

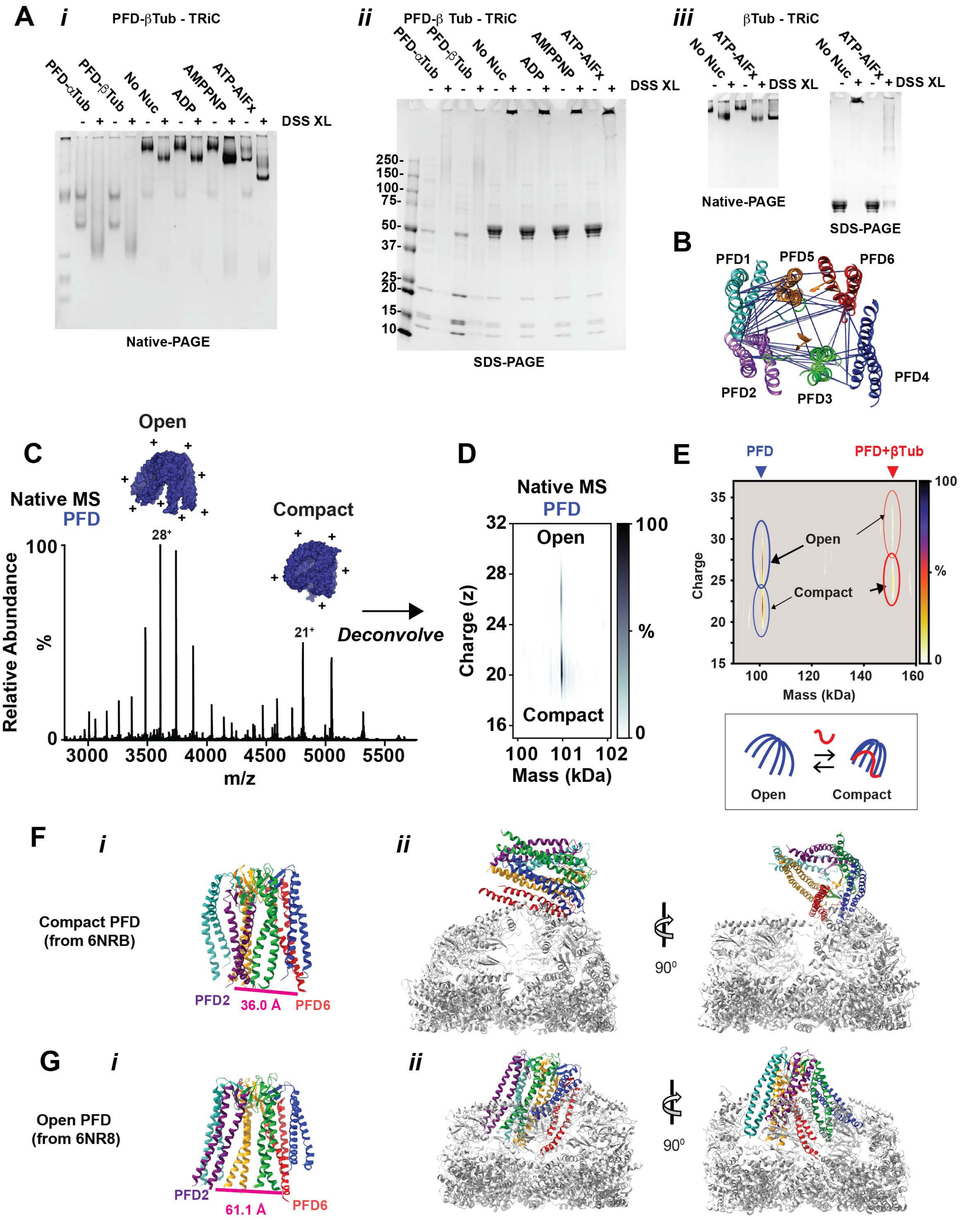
A
i
PFD-βTub - TRiC
PFD-αTub
PFD-βTub
No Nuc
ADP
AMPPNP
ATP-AlFx
DSS XL
Native-PAGE
ii
PFD-β Tub - TRiC
250-
150-
100-
75-
50-
37-
25-
20-
15-
10-
SDS-PAGE
iii
βTub - TRiC
Native-PAGE
SDS-PAGE
B
PFD1
PFD5
PFD6
PFD4
PFD2
PFD3
C
Open
Native MS
PFD
Compact
28+
21+
Relative Abundance
%
Deconvolve
m/z
D
Native MS
PFD
Open
Compact
Charge (z)
Mass (kDa)
E
PFD
PFD+βTub
Open
Compact
Charge
Mass (kDa)
%
Open
Compact
F
i
ii
Compact PFD
(from 6NRB)
PFD2
36.0 Å
PFD6
90⁰
G
i
ii
Open PFD
(from 6NR8)
PFD2
61.1 Å
PFD6
90⁰

**Figure S2. Native MS and XL analysis for PFD and PFD-βTub; Related to figure 2.**
(A) Native PAGE (i, iii) and SDS-PAGE (ii, iii) from XLing of PFD-Tub, PFD-Tub-TRiC, and Tub-TRiC samples in indicated nucleotide condition. In native gel XLed samples run as single complexes, while in SDS-PAGE XLed samples migrate as multi-protein species due to interXLing. (B) Bottom view of XLs mapped onto PFD, note extensive XLs between PFD2 and PFD6. (C) Representative native mass spectrum of PFD without βTub. Two charge state distributions can be inferred to correspond to conformations that are relatively open and compact, represented by cartoons with more or fewer charges (+). The most abundant charge state of each conformation is labeled. (D) Mass to charge plot of deconvolved data in (C). A single mass, PFD, gives rise to both peak distributions of charge. (E) Alternative representation of the native mass spectrometry data in main figure 2F, with charge plotted on the vertical axis and intensity as a color scale (right). Open and compact states of PFD and PFD-βTub are outlined in blue and red respectively, with outline widths proportional to relative abundances. These data show a shift in conformational equilibrium upon βTub binding. (F,G) PFD structures alone (i) and with the top ring of TRiC (ii) from the least engaged (F) PDB:6NRB and most engaged (G) PDB:6NR8 PFD-TRiC structures. The distance between the terminally resolved residues of PFD2 (Ile 124) and PFD6 (Glu 114) are indicated in magenta.

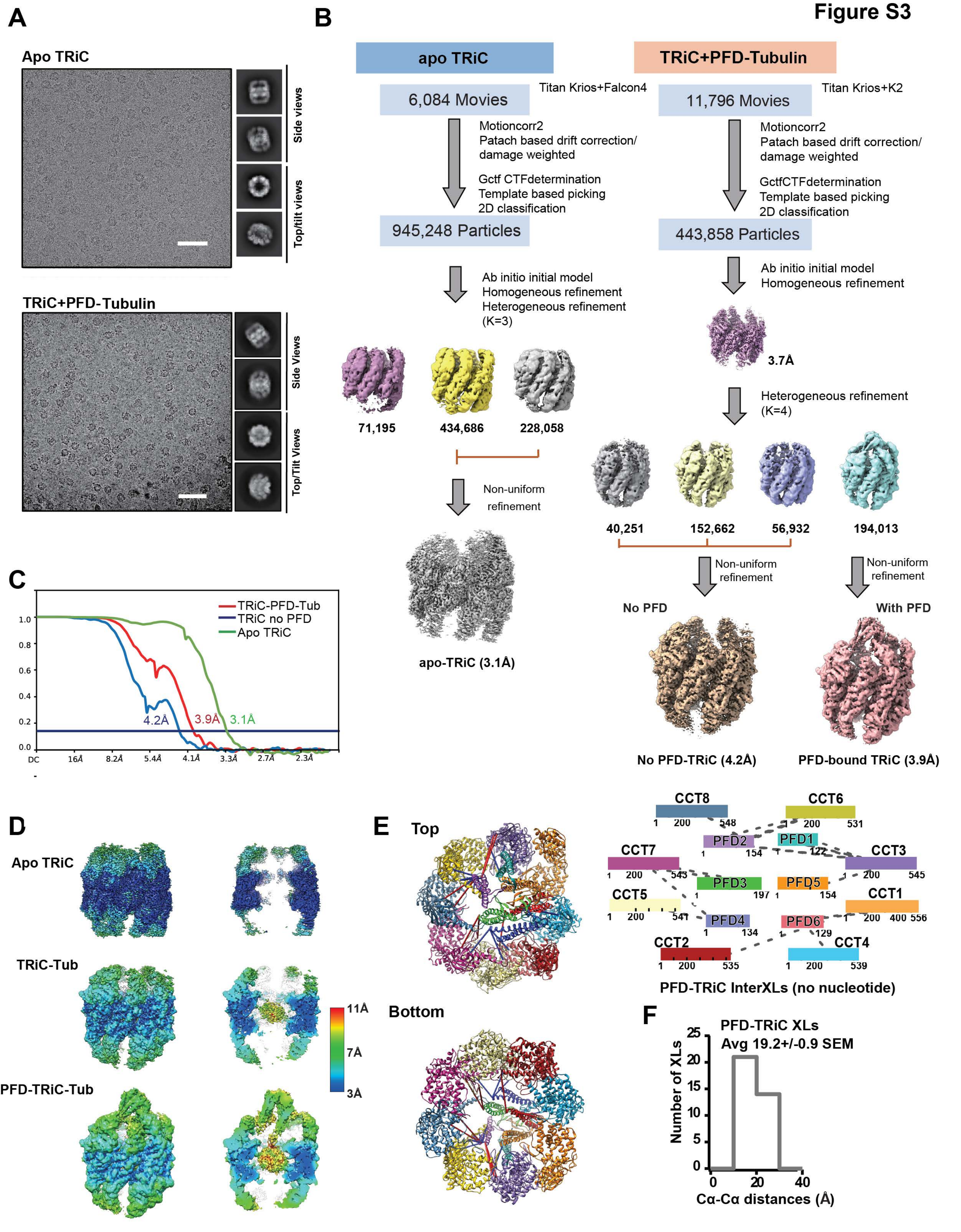
Figure S3
A
Apo TRiC
Side views
Top/tilt views
TRiC+PFD-Tubulin
Side Views
Top/Tilt Views
B
apo TRiC
TRiC+PFD-Tubulin
6,084 Movies
Titan Krios+Falcon4
11,796 Movies
Titan Krios+K2
Motioncorr2
Patach based drift correction/
damage weighted
Gctf CTFdetermination
Template based picking
2D classification
GctfCTFdetermination
945,248 Particles
443,858 Particles
Ab initio initial model
Homogeneous refinement
Heterogeneous refinement
(K=3)
71,195
434,686
228,058
Non-uniform
refinement
apo-TRiC (3.1Å)
3.7Å
Heterogeneous refinement
(K=4)
40,251
152,662
56,932
194,013
No PFD
With PFD
No PFD-TRiC (4.2Å)
PFD-bound TRiC (3.9Å)
C
TRiC-PFD-Tub
TRiC no PFD
Apo TRiC
4.2Å
3.9Å
3.1Å
1.0
0.8
0.6
0.4
0.2
0.0
DC
16Å
8.2Å
5.4Å
4.1Å
3.3Å
2.7Å
2.3Å
D
Apo TRiC
TRiC-Tub
PFD-TRiC-Tub
11Å
7Å
3Å
E
Top
Bottom
CCT8
CCT6
PFD2
PFD1
CCT7
CCT3
PFD3
PFD5
CCT5
CCT1
PFD4
PFD6
CCT2
CCT4
PFD-TRiC InterXLs (no nucleotide)
F
PFD-TRiC XLs
Avg 19.2+/-0.9 SEM
Number of XLs
Cα-Cα distances (Å)

**Figure S3: CryoEM workflow and XL analysis for PFD-βTub-TRiC; Related to figure 3.**

(A) Representative cryoEM micrograph and 2D averages of apo-TRiC and the complex of PFD-βTub-TRiC in the open conformation. (B) Image processing workflow of apo-TRiC and PFD-βTub-TRiC and Gold standard FSC curve (C). (D) Local resolution estimation of three reconstructed maps: apo-TRiC, TRiC-βTub (PFD unbound), PFD-TRiC-βTub. (E) Crosslink mapping on PFD-βTub bound to TRiC (F) Histogram of shortest measured intra/interring Cα -Cα distances for identified XLs between PFD and TRiC.

Figure S4

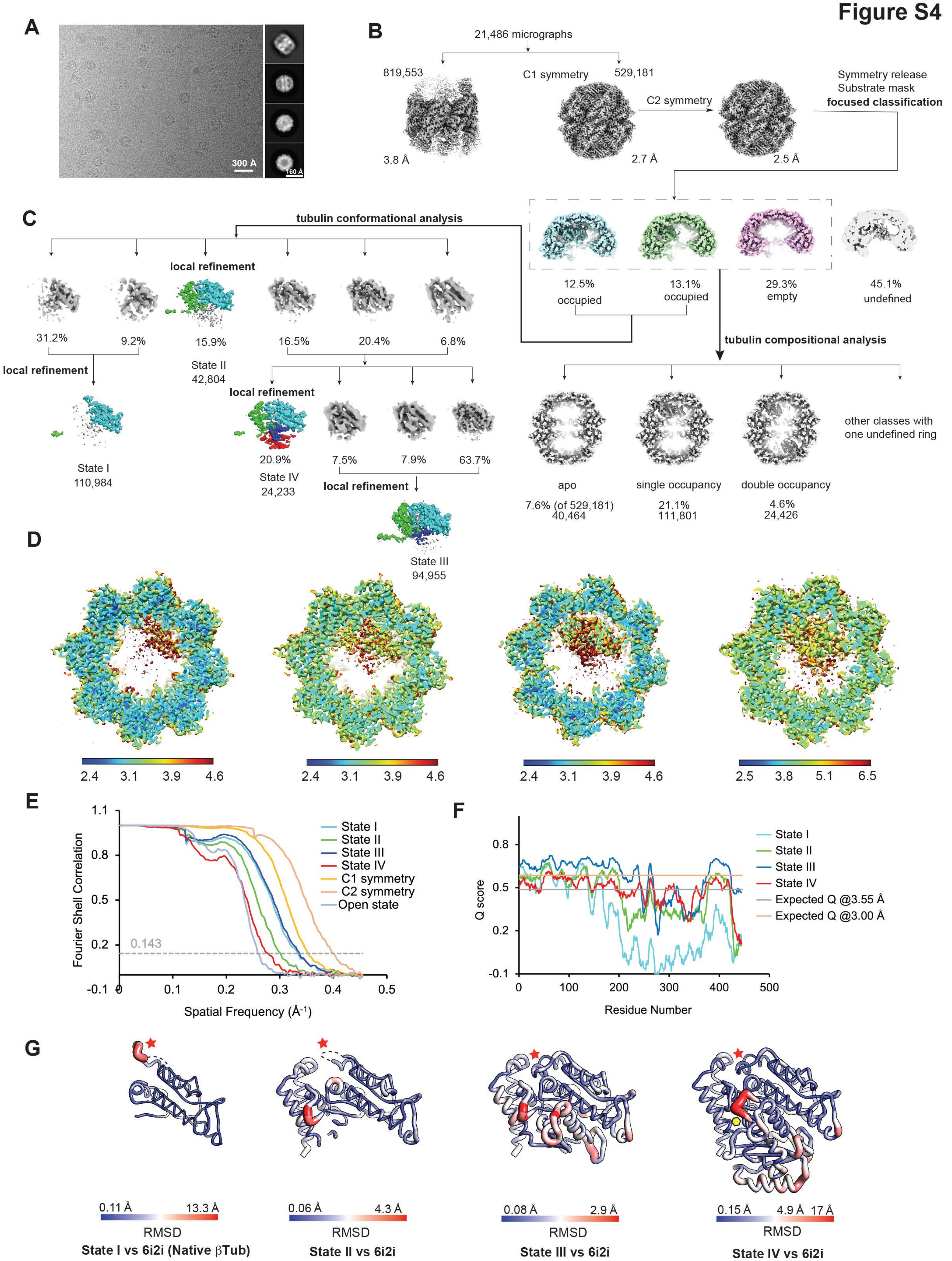
A
300 Å
160 Å
B
21,486 micrographs
819,553
C1 symmetry
529,181
C2 symmetry
Symmetry release
Substrate mask
focused classification
3.8 Å
2.7 Å
2.5 Å
C
tubulin conformational analysis
local refinement
12.5% occupied
13.1% occupied
29.3% empty
45.1% undefined
31.2%
9.2%
15.9%
16.5%
20.4%
6.8%
State II
42,804
tubulin compositional analysis
local refinement
local refinement
State I
110,984
20.9%
7.5%
7.9%
63.7%
State IV
24,233
local refinement
other classes with one undefined ring
apo
single occupancy
double occupancy
7.6% (of 529,181)
40,464
21.1%
111,801
4.6%
24,426
State III
94,955
D
2.4 3.1 3.9 4.6
2.4 3.1 3.9 4.6
2.4 3.1 3.9 4.6
2.5 3.8 5.1 6.5
E
Fourier Shell Correlation
Spatial Frequency (Å-1)
0.143
State I
State II
State III
State IV
C1 symmetry
C2 symmetry
Open state
F
Q score
Residue Number
State I
State II
State III
State IV
Expected Q @3.55 Å
Expected Q @3.00 Å
G
0.11 Å 13.3 Å
RMSD
State I vs 6i2i (Native βTub)
0.06 Å 4.3 Å
RMSD
State II vs 6i2i
0.08 Å 2.9 Å
RMSD
State III vs 6i2i
0.15 Å 4.9 Å 17 Å
RMSD
State IV vs 6i2i

**Figure S4. CryoEM workflow and analysis for TRiC-βTub; Related to figure 4.**

(A) Representative micrograph and 2D class averages of TRiC-βTub with ATP-AlFx. (B) Processing workflow of TRiC-βTub with ATP-AlFx and (C) 3D classification strategy for focusing the density inner chamber of TRiC. Our compositional heterogeneity analysis identifies different βTub occupancies for TRiC, including no substrate bound (*apo* state), and single or double ring occupancy. Further analysis identified four progressively folded tubulin conformations inside the chamber: State I (41% of the rings); State II (16% of the rings); State III (34% of the rings) and State IV (9% of the rings) corresponding to folded tubulin. (D) Local resolution map estimation of βTub intermediate states. (E) FSC curve of reconstructed maps showing resolution according to the gold standard FSC (0.143 criterion). (F) Q-score plots of the βTub intermediate states shown with the expected Q-score according to 3.55 and 3.00 Å. (G) C-alpha backbone RMSD per residue between intermediate state and native βTub (PDB: 6I2I). Large variation indicated at T3 loop (red star) and N loop (yellow hexagon)

Figure S5

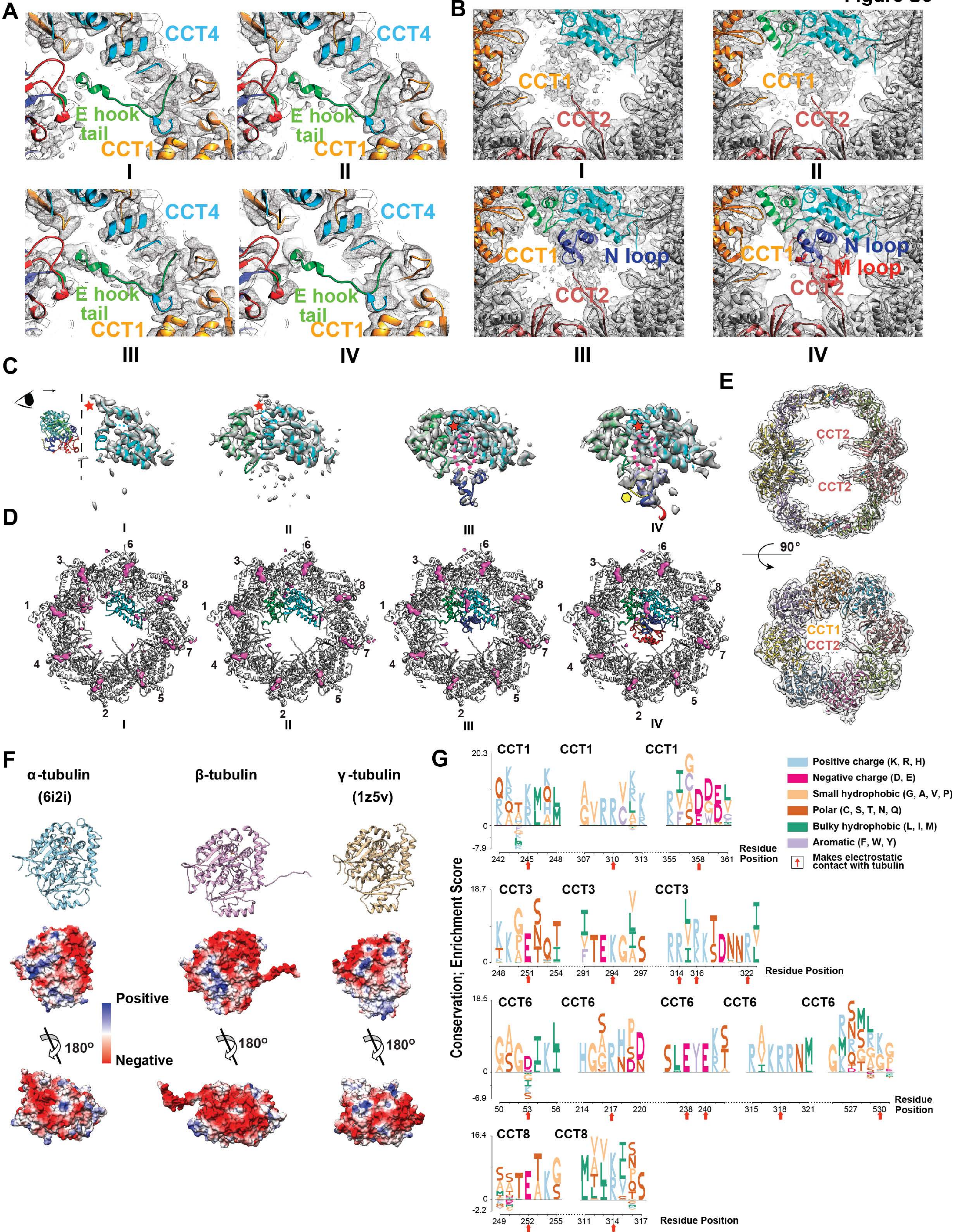
A
CCT4
E hook tail
CCT1
I
II
III
IV
B
CCT1
CCT2
N loop
M loop
C
D
1
2
3
4
5
6
7
8
E
CCT2
CCT1
90°
F
α-tubulin
(6i2i)
β-tubulin
γ-tubulin
(1z5v)
Positive
Negative
180°
G
Conservation; Enrichment Score
CCT1
CCT3
CCT6
CCT8
Residue Position
Positive charge (K, R, H)
Negative charge (D, E)
Small hydrophobic (G, A, V, P)
Polar (C, S, T, N, Q)
Bulky hydrophobic (L, I, M)
Aromatic (F, W, Y)
Makes electrostatic contact with tubulin

**Figure S5. TRiC chamber spatially orients and restrains βTub; Related to figure 5.**

(A) CryoEM density of βTub E hook tail in the CCT1/4 pocket for the different folding states (I-IV). (B) CryoEM density of CCT2 C terminal tail connected to βTub for the different folding states (I-IV). (C) CryoEM density of the βTub at the GTP binding pocket: T3 loop (red star), N loop (yellow hexagon), nucleotide (magenta eclipse). (D) Difference map analysis of the nucleotide occupancy in βTub GTP binding pocket in comparison to CCT ATP binding pockets. The difference maps are displayed with +3σ. (E) Slice side and top view of CCT1/2 C terminal tail contacts in *apo* closed TRiC. (F) Electrostatic surface display of human αTub (PDB: 6i2i), βTub (PDB: ours) and γTub (1z5v), respectively. (I) Logo plot of the conservation of CCT residues. The residues making direct contact with βTub are indicated by red arrows

### Figure S6

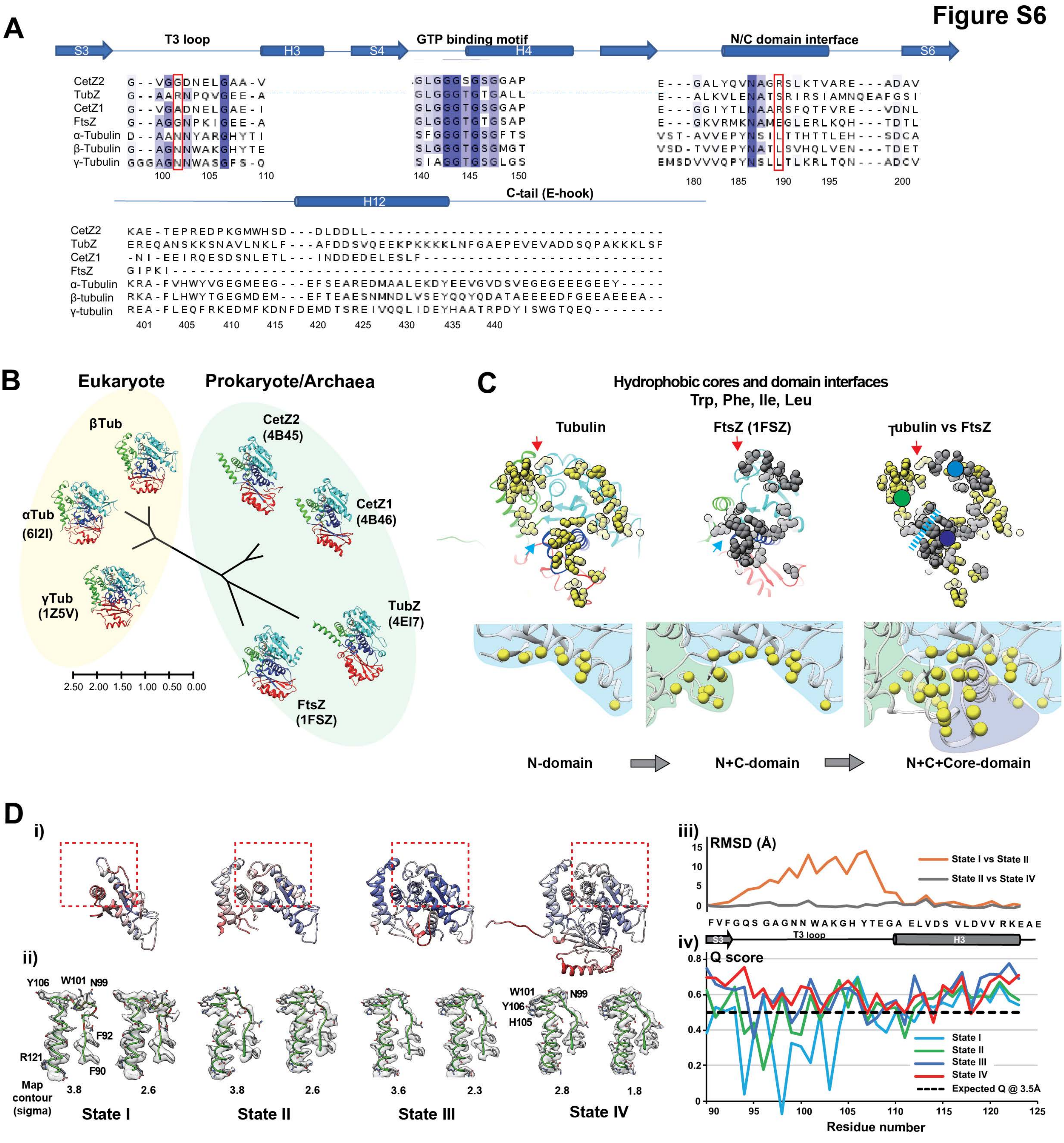


## E

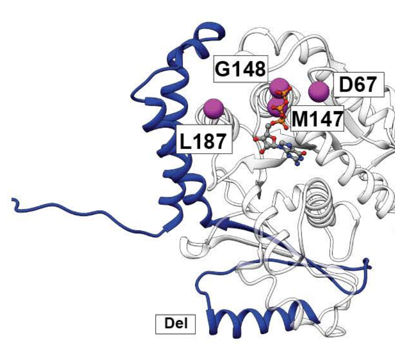


| Mutant | Results | Category | Location | Reference |
|---|---|---|---|---|
| M147K, G148T | Interferes the ability to fold | GTP binding pocket | H4 helix | Juan et al., 1996 |
| D67N | More susceptible to proteolysis | GTP binding pocket | S2 strand | Farr and Sternlicht, 1992 |
| L187R | Not released from TRiC | N/C domain interface | H5 helix | Wang et al., 2006 |
| Deletion | Not released from TRiC | C truncation | Residue 325-444 | |

**Figure S6. Tubulin coevolved with TRiC for directed folding process; Related to figure 6.**

(A) Sequence alignment for selected areas of human α, β, γ tubulins and archaeal/prokaryotic homologues (CetZ1: H. Volcanii CetZ2: H. Volcanii, FtsZ: M. jannaschii and TubZ: B.cereus). T3 loop, GTP binding motif, N/C-domain interface and C-domain including H12 and E-hook are represented. Red box indicates conserved N99 and L189 in T3 loop and N/C-domain interface in human tubulins. (B) Phylogenetic tree of tubulins and their structures. Structures are colored in domain (N-domain: cyan, C-domain: green, Core domain: blue, M-domain: red). The scale bar represents the number of amino acids substitutions per site. (C) The hydrophobic core and domain interfaces of βTub and FtsZ with hyrdrophobic residues (Trp, Phe, Ile, Leu) highlighted with yellow (βTub) and gray (FtsZ) balls. Red arrow indicates the N/C-domain interface. Blue arrows and dashed line indicate the C/Core-domain interface. Colored circles represent hydrophobic core of each domain. The progressive assembly of the hydrophobic cores for N-, C- and Core domain of βTub (bottom panels). (D) Model B factor distribution of each state highlights the N-domain with T3 loop in red dashed box (i) and corresponding cryo-EM density maps with fitted model are presented (ii). Per residue RMSD plots of T3 loop among four states (iii) and Q-score plot of T3 loop (iv). Dashed black line indicates average Q score expected at 3.5 Å resolution. (E) βTub model with point mutations at the GTP binding pocket indicated as balls on residues that have folding defects. Their locations and effects are summarized (Farr and Sternlicht, 1992; Wang et al., 2006; Zabala et al., 1996).

**Legend to Supplemental Movies**

**Movie S1. Cryo-EM density of the four βTub folding states in closed TRiC chamber; Related to figure 4.** This movie demonstrates the local density of three βTub intermediates and fully folded states. The model of the folded βTub domain is fitted into each density map and displayed. GTP is fitted into the GTP binding pocket as an example to illustrate the observed extra density in State III and IV.

**Movie S2. Asymmetric charged surface of TRiC closed chamber; Related to figure 5.** This movie illustrates that closed TRiC has an asymmetrically charged inner chamber wall.

**Movie S3. Electrostatic interaction between βTub N-, C-domain and the TRiC chamber wall; Related to figure 5.** This movie illustrates βTub N-, C-domain interaction with the CCT1/3/6/8 by complementary charged surface.

**Movie S4. Description of the E hook tail of βTub and CCT2/6 C-terminal tail in the βTub folding process; Related to figure 5.** This movie illustrates that the E hook tail of βTub inserts into the CCT1/4 pocket, CCT6 termini contacts the GTP binding pocket and CCT2 C-termini interacts the Core/M-domain at the N/M loop.

**Movie S5. Inter-domain hydrophobic burial in the βTub folding process and continuous hydrophobic framing of the growing β-sheet in the folding process; Related to figure 6.** The first part of the movie illustrates that TRiC orients hydrophobic surface of βTub N-, C-domain to face the chamber center and allows the subsequent hydrophobic burial with Core helix and M-domain folding. The second part of movie demonstrates the progressive β-sheet building along with continuously significant hydrophobic framing from helices as βTub folds.

**Movie S6. Conformational change of T3 loop from State I to the other states; Related to figure 6.** The movie shows the model trajectory from state I to the other states highlighting a conformational change of T3 loop in βTub.

**Table S1. CryoEM image collection, map reconstruction, and model refinement**

| | Apo-TRiC | PFD-βTub-TRiC | TRiC-βTub State I | TRiC-βTub State II | TRiC-βTub State III | TRiC-βTub State IV |
|---|---|---|---|---|---|---|
| **CryoEM image collection and map processing** | | | | | | |
| Voltage (kV) | 300 | 300 | 300 | 300 | 300 | 300 |
| Total electron exposure ($e^{-}/Å^2$) | 37 | 36 | 37 | 37 | 37 | 37 |
| Defocus range (μm) | -0,5~-2.5 | -0.5~-3.5 | -0.5~-3.5 | -0.5~-3.5 | -0.5~-3.5 | -0.5~-3.5 |
| Pixel size (Å) | 1.0 | 1.02 | 1.1 | 1.1 | 1.1 | 1.1 |
| Symmetry imposed | C1 | C1 | C1 | C1 | C1 | C1 |
| Particle images (no.) | 662,744 | 194,013 | 110,984 | 42,804 | 94,955 | 24,233 |
| Map resolution (Å)<br>FSC threshold (0.143) | 3.1 | 3.9 | 3.0 | 3.3 | 2.9 | 3.6 |
| EMDB ID | 32822 | 32823 | 26089 | 26120 | 26123 | 26131 |
| **Model refinement** | | | | | | |
| Initial model used (PDB code) | NA | 6NR8 | 7LUM, 6I2I | 7LUM, 6I2I | 7LUM, 6I2I | 7LUM, 6I2I |
| Model composition<br>Non-hydrogen atoms<br>Protein residues<br>Ligands | | <br>67039<br>8727<br>ADP:4 | <br>33714<br>4377<br>MG: 8<br>ADP: 8<br>$AF_3$: 8<br>$H_2O$: 8 | <br>34563<br>4481<br>MG: 8<br>ADP: 8<br>$AF_3$: 8<br>$H_2O$: 8 | <br>35180<br>4561<br>MG: 8<br>ADP: 8<br>$AF_3$: 8<br>$H_2O$: 8 | <br>35877<br>4650<br>MG: 8<br>ADP: 8<br>$AF_3$: 8<br>$H_2O$: 8 |
| R.m.s. deviations<br>Bond lengths (Å)<br>Bond angles (°) | | <br>0.003<br>0.790 | <br>0.014<br>1.137 | <br>0.012<br>1.233 | <br>0.010<br>1.049 | <br>0.006<br>0.767 |
| Validation<br>MolProbity score<br>Clashscore<br>Rotamer outliers (%) | | <br>2.12<br>18.09<br>0.08 | <br>1.6<br>4.7<br>0.08 | <br>1.61<br>4.54<br>0.32 | <br>1.57<br>4.52<br>0.00 | <br>1.65<br>6.22<br>0.13 |
| Ramachandran plot<br>Favored (%)<br>Allowed (%)<br>Outliers (%) | | <br>94.66<br>5.25<br>0.09 | <br>94.83<br>5.03<br>0.14 | <br>94.39<br>5.45<br>0.16 | <br>95.02<br>4.78<br>0.20 | <br>95.62<br>4.30<br>0.09 |
| PDB ID | | 7WU7 | 7TRG | 7TTN | 7TTT | 7TUB |

**Table S2: Summary of XL-MS data (XLS file)**

**CONTACT FOR REAGENT AND RESOURCE SHARING**

**DETAILED EXPERIMENTAL PROCEDURES**

##### Cloning

*pFB-αTubulin and pFB-βTubulin:* Human Beta-Tubulin (TBB5) and Alpha-Tubulin (TUBA1B) were cloned into the pFB-Dual vector using conventional methods. Briefly, βTubulin was amplified using 5’ primer TTT TAG CGG CCG CCA AAA CAT GAG GGA AAT CGT GCA CAT CC and 3’ primer ATT TTT CTA GAT TAT CAG GCC TCC TCT TCG GCC TCC, and αTubulin was amplified using 5’ primer TTT TAG CGG CCG CCA AAA CAT GCG TGA GTG CAT CTC CAT CC and 3’ primer ATT TTT CTA GAT TAT CAG TAT TCC TCT CCT TCT TCC TCA CC from the plasmids pTS3067 and pTS4119 respectively. Amplified products and pFB-Dual vector were combined after digestion with NotI and XbaI, and sequence of isolated clones was verified.

*TBCA and TBCB:* TBCA was cloned into pST39 and TBCB was cloned into pFB-Dual vectors using conventional methods. Briefly, TBCA was amplified using 5’ primer AGT TTC TAG ATT TGT TTA ACT TTA AGA AGG AGA TAT ACA TAT GGC CGA TCC TCG CGT GAG and 3’ primer AAT TAG GAT CCT CAT TAG TGA TGG TGA TGA TGG TGG GCT TCT AAC TTC ACT G and cloned into pST39 using XbaI and BamHI. TBCB was first amplified using 5’ primer AGT TTC TAG ATT TGT TTA ACT TTA AGA AGG AGA TAT ACA TAT GGA GGT GAC GGG GGT GTC and 3’ primer (with 6XHis) AAT TAG GAT CCT CAT TAG TGA TGG TGA TGA TGG TGA CTG CCT ATC TCG TCC AAC CCG TAG TCC and cloned into pST39 with XbaI and BamHI. To move TBCB into the pFB-Dual vector for insect cell expression TBCB was PCR amplified out of pST39 using 5’ TAA TTC TCG AGC TAT AAA TAT GGA GGT GAC GGG GGT GTC G and 3’ TAA TTG GTA CCT CAT TAG TGA TGG TGA TGA TGG TGA CTG CC and inserted into pFB-Dual using XhoI and KpnI.

##### Protein expression and purification

*TRiC alone*. High Five insect cells (1 L) were co-infected with rBVs encoding his-tagged TRiC with CCT1 tagged with GFP. Cells were incubated at 27°C for ~ 72 h, pelleted by centrifugation at 500 x g for 10 min, and resuspended in TRiC lysis buffer (100 mM HEPES pH 7.4, 50 mM NaCl, 20 mM imidazole, 10 % glycerol, 5 mM PMSF) supplemented with benzonase (Sigma-

Aldrich-Aldrich, E1014) (1,000 units) and a protease inhibitor cocktail [Roche]). Cells were lysed using dounce homogenization and debris was cleared by ultracentrifugation at 50,000 x g at 4 °C for 40 min. Cleared supernatant was passed over nickel resin and washed with column wash buffer (50 mM HEPES pH 7.4, 50 mM NaCl, 5 mM $MgCl_2$ 20 mM imidazole, 10 % glycerol) with an additional 250 mM NaCl, column wash buffer + 1 mM ATP, column wash buffer + an additional 500 mM NaCl, and finally column wash buffer alone. Nickel-bound protein was eluted with elution buffer (50 mM HEPES pH 7.4, 50 mM NaCl, 5 mM $MgCl_2$ 400 mM imidazole, 10 % glycerol). Protein-containing fractions were pooled and passed over a heparin column equilibrated with MQA buffer (50 mM HEPES pH 7.4, 50 mM NaCl, 5 mM $MgCl_2$, 0.5 mM EDTA, 1 mM DTT, 10 % glycerol). Protein was eluted with a linear gradient of 20 % to 100 % MQB buffer (50 mM HEPES pH 7.4, 1 M NaCl, 5 mM $MgCl_2$, 0.5 mM EDTA, 1 mM DTT, 10 % glycerol). TRiC-containing fractions were pooled and diluted with MQ buffer (50 mM HEPES pH 7.4, 5 mM $MgCl_2$, 1 mM DTT, 10 % glycerol) to remove excess NaCl. Pooled TRiC was loaded onto a MonoQ ion exchange column and eluted with a 200 ml linear gradient of 0 % to 100 % MQB. TRiC containing fractions were pooled, concentrated with a 100 kDa MWCO Centricon device, and passed over a Superose 6 size exclusion column equilibrated with MQA. TRiC containing fractions were pooled, concentrated, and snap-frozen in liquid nitrogen for long-term storage.

*TRiC-βTubulin purification:* High Five insect cells (1 L) were co-infected with rBVs encoding his-tagged TRiC and the Human βTubulin protein. Cells were incubated at 27°C for ~ 72 h, pelleted by centrifugation at 500 x g for 10 min, and resuspended in TRiC lysis buffer (100 mM HEPES pH 7.4, 50 mM NaCl, 20 mM imidazole, 10 % glycerol, 5 mM PMSF) supplemented with benzonase (Sigma-Aldrich-Aldrich, E1014) (1,000 units) and a protease inhibitor cocktail [Roche]). Cells were lysed using dounce homogenization and debris was cleared by ultracentrifugation at 50,000 x g at 4 °C for 40 min. Cleared supernatant was passed over nickel resin and washed with column wash buffer (50 mM HEPES pH 7.4, 50 mM NaCl, 20 mM imidazole, 10 % glycerol). Nickel-bound protein was eluted with column wash buffer containing 400 mM imidazole. Protein-containing fractions were pooled and passed over a heparin column equilibrated with MQA buffer (50 mM HEPES pH 7.4, 50 mM NaCl, 5 mM $MgCl_2$, 0.5 mM EDTA, 1 mM DTT, 10 % glycerol). Protein was eluted with a linear gradient of 20 % to 100 % MQB buffer (50 mM HEPES pH 7.4, 1 M NaCl, 5 mM $MgCl_2$, 0.5 mM EDTA, 1 mM DTT, 10 % glycerol). TRiC-containing fractions were pooled and diluted with MQ buffer (50 mM HEPES pH 7.4, 5 mM $MgCl_2$, 1 mM DTT, 10 % glycerol) to remove excess NaCl. Pooled TRiC was loaded onto a MonoQ ion exchange column and eluted with a 200 ml linear gradient of 0 % to 100 % MQB. Fractions were assayed for the presence of βTub and TRiC, pooled, concentrated with a 100 kDa MWCO Centricon device, and passed over a Superose 6 size exclusion column equilibrated with MQA. TRiC containing fractions were pooled, concentrated, and snap-frozen in liquid nitrogen for long-term storage. The integrity of TRiC and βTub occupancy were confirmed by SDS-PAGE, native-PAGE, Coomassie staining, and immunoblotting with a rabbit CCT5-specific antibody (Abcam ab129016) and a mouse βTub specific monoclonal antibody (Proteintech 66240-1-Ig).

*Human PFD:* Human PFD was expressed and purified as described previously (Gestaut et al., 2019). All subunits were co-expressed using baculovirus in High Five insect cells. The PFD complex was isolated by affinity (Nickel-NTA), anion exchange (MonoQ 10/100), and gel filtration (Superdex 200) chromatography. At each step, fractions containing PFD were identified by SDS-PAGE. Final protein was concentrated to ~100 µM using an Amicon Ultra 30 kDa MWCO, and 50 % glycerol was added to yield a final concentration of 10 %. Protein was aliquoted, snap-frozen, and stored at -80 °C.

*Human PFD + αTubulin, βTubulin:* Human PFD was co-expressed with human αTubulin or βTubulin in High Five insect cells and purified as described above without the anion exchange step. All subunits were coexpressed using baculovirus in High Five insect cells. The PFD complex was isolated by affinity (NickelNTA) and gel filtration (Superdex 200) chromatography. At each step, fractions containing PFD and Tubulin were identified by SDS-PAGE. Final protein was concentrated to ~ 20 µM using an Amicon Ultra 30 kDa MWCO, and 50 % glycerol was added to yield a final concentration of 10 %. Protein was aliquoted, snap-frozen, and stored at -80 °C.

*TBCA:* TBCA was expressed in 4 L of BL21 Rosetta2 pLysS cells (Novagen) O/N at 16 °C. Cells were lysed in 50 ml of column buffer (300 mM NaCl, 50 mM HEPES pH 7.4) with benzonase, fresh PMSF, 10 mM imidazole and Complete protease inhibtiors, EDTA free. Cells were lysed 2X using an emulsiflex (Avestin), and lysate was cleared at 20,000 x g for 30 min. Protein was affinity purified in a 10 ml Ni column equilibrated with column buffer and eluted with column buffer + 400 mM imidazole after a wash of ~300 ml column buffer + 10 mM imidazole. Protein was concentrated using an Amicon ultra 3 kD MWCO device to less than 2 ml and run over an Superdex 75 SEC column equilibrated with column buffer + 1 mM DTT. Protein was concentrated using an Amicon ultra 3 kD MWCO device, glycerol added to 5 %, aliquoted, snap-frozen, and stored at -80 °C.

<u>*TBCB:*</u> TBCB was expressed in Hi-5 cells adapted to ESF media by infection with 15 ml of P2 virus into 1 L of cells at ~ 2 x $10^6$ cells/ml. Cells were harvested 3 days after infection. Cells were lysed in column buffer (300 mM NaCl, 50 mM HEPES pH 7.4) with Benzonase, fresh PMSF, 20 mM imidazole and Complete protease inhibitors, EDTA free. Cell pellet was resuspended in ~90 ml lysis buffer and lysed using dounce homogenization. Lysate was cleared at 40,000 x g for 45 min. Protein was affinity purified in a 25 ml Ni column equilibrated with column buffer and eluted with column buffer + 400 mM imidazole after washes of ~75 ml column buffer + 15 mM imidazole, 50 ml column buffer + 500 mM NaCl + 15 mM imidazole, 50 ml column buffer + 15 mM imidazole. Protein was concentrated using an Amicon ultra 3 kD MWCO to less than 2 ml and run over an Superdex 200 SEC column equilibrated with 200 mM NaCl, 50 mM HEPES pH 7.4, 1 mM TCEP. Protein was concentrated using an Amicon ultra 3 kD MWCO device, glycerol added to 5 %, aliquoted, snap-frozen, and stored at -80 °C.

<u>*mmCPN:*</u> MmCpn was expressed off pET21a containing E. coli Rosetta DE3 pLys (Novagen) and 2 L were grown to 0.3 OD and induced at 17 °C O/N. Cells were harvested by pelleting at 4500 x g for 10 min, and washed once with cold PBS (137 mM NaCl, 2.7 mM KCl, 8 mM $Na_2HPO_4$, and 2 mM $KH_2PO_4$). Cells were pelleted again at 4500 x g for 10 min, and resuspended in MQA (50 mM HEPES pH 7.4, 50 mM NaCl, 10 % glycerol, 5 mM PMSF) supplemented with 1 mM DTT, benzonase (Sigma-Aldrich-Aldrich, E1014) (1,000 units) and a protease inhibitor cocktail [Roche]). Cells were lysed 3X using an emulsiflex (Avestin). Lysate was cleared at 20,000 x g for 30 min. Ammonium sulfate was slowly added to the cleared lysate to 55% and left on mutator for 10 min. Lysate was cleared at 20,000 x g for 20 min, and supernatant was dialyzed O/N in MQA. Dialzyed lysate loaded onto an equilibrated CaptoQ (Cytiva) column (~22 ml resin), column was washed until UV abs baselined, and protein was eluted with a 150 ml gradient from MQA to MQB (50 mM HEPES pH 7.4, 1 M NaCl, 10 % glycerol, 5 mM PMSF). Fractions containing mmCPN were identified by SDS-PAGE, combined (12 mls) and diluted to 50 mls to lower NaCl concentration. Diluted protein was loaded onto a HiPrep Heparin FF 20 ml column (Cytiva) pre-equilibrated with 20% MQB, washed with 25% MQB, and eluted with a 160 ml gradient to 100% MQB. Fractions containing mmCPN were identified by SDS-PAGE, combined, and concentrated to ~ 1 ml using an Amicon Ultra 30 kD MWCO (4ml) (EMD Millipore). Protein was further purified using SEC on a Superose 6 equilibrated with MQA. Fractions containing mmCPN were identified by SDS-PAGE, combined, concentrated to ~ 300 μl using an Amicon Ultra 30 kD MWCO, aliquoted, snap frozen and stored at -80 °C.

**Native and SDS PAGE analysis**

_Native and SDS gel analysis for Prefoldin-tubulin:_ Prefoldin samples were diluted to 5 μM. For native gels 7.5 μl of sample was run out on 4-16 % Native-PAGE gels. For SDS-PAGE, 8 μl of 4XPSB (200 mM Tris pH 6.8, 8% SDS, 40% glycerol, 20% β-mercaptoethanol, bromophenol blue to color) was added to 24 μl of sample, and 10 μl of each sample was separated in 15 % SDS-PAGE gels. Both gels were run in duplicate with one used for Coomassie staining and the other for immunoblot.

_Native and SDS gel analysis for TRiC-tubulin:_ TRiC samples were diluted to 0.5 μg/μl and bovine Tubulin to ~0.05 μg/μl for each Tubulin. For native gels, 5 μl of TRiC samples was separated. SDS-PAGE samples were made by adding 4XPSB to 1X final concentration, 6 μl of TRiC samples and 1, 2, 4 and 8 μl of Tubulin were separated.

**Tubulin folding assays**

βTub folding assay was performed in folding buffer (20 mM Tris-HCl pH 7.4, 50 mM KCl, 10 mM $MgCl_2$ and 0.2 mM $CaCl_2$) (Balchin et al., 2018) with fresh TCEP added to 1 mM. PFD-βTub was prepared to a final concentration of 1 μM, TRiC to 0.1 μM, and TBCA to 1 μM. Samples were incubated in the presence/absence of 0.9 mM ATP, 1 mM GTP as indicated for 1 h. Folding assay samples were separated by native page and βTub was detected by immunoblot with a monoclonal mouse antibody (Proteintech 66240-1-Ig). αTub folding was performed in folding buffer with an additional 50 mM KCl and fresh TCEP added to 1 mM. PFD-αTub was prepared to a final concentration of 1 μM, TRiC to 0.25 μM, and TBCB to 1 μM. Samples were incubated in the presence/absence of 0.9 mM ATP, 1 mM GTP as indicated for 1 h at 37 C. Folding assay samples were separated by native page and αTub was detected by immunoblot with a monoclonal mouse antibody (Sigma-Aldrich T5168).

**Proteinase K assays**

Prefoldin-Tubulin, Bovine Tubulin, TRiC, and mmCPN were brought to a final concentration of 1 μM in 1XATPase buffer (50 mM HEPES, 50 mM KCl, 5 mM $MgCl_2$, 10 % glycerol, 1 mM DTT). Samples were brought to 37 °C and 1 mM ATP/GTP was added to the Tubulin dimer control, PFD-Tubulin sample, TRiC-PFD-Tubulin sample (+ATP/GTP), and TRiC-PFD-Tubulin sample

(+ATP/GTP-AlFx). After 5 min, NaF to 1 mM, and $AlNO_3$ to 6 mM was added to the TRiC-PFD-Tubulin sample (+ATP/GTP-AlFx). After 30 min, samples were brought to room temperature. 15 µl was removed from each sample for zero-minute time points, and digestion was initiated by addition of proteinase K to a final concentration of 2 ng/µl. Timepoints were taken at 2, 4, 8 and 16 min by adding 15 µl sample to 2 µl of PMSF, followed by 6 µl 4XPSB. Samples were analyzed by SDS-PAGE followed by immunoblot with indicated antibodies. For assays with denatured Tubulin, chaperones were diluted to 0.25 µM. Bovine Tubulin at a concentration of 12.5 µM was denatured in 6 M GuHCl, 100 mM HEPES pH 7.4, 1 mM DTT, and chaperone mixes were used to dilute denatured Tubulin 1:100 for a final concentration of 0.125 µM. Aggregated Tubulin was pelleted by centrifugation at 21 K*G for 10 min. Supernatant was removed and PK assay was carried out as described as above with 14 µl timepoints. Tubulin binding by chaperones was assayed by running 10 µl samples in Native PAGE. For western blots, βTub was probed for with a monoclonal mouse antibody towards the N-terminus (Proteintech 66240-1-Ig), a polyclonal rabbit antibody (abcam ab6046) or a C-terminal epitope specific AA2 antibody (Millipore sigma 05-661), and αTub with a mouse monoclonal antibody (Sigma-Aldrich T5168).

**Crosslinking-Mass Spectrometry**

For hPFD-βTub with hTRiC samples, TRiC was brought to 1 µM, and PFD to 2 µM, hPFD-α/βTub alone was prepared at 5 µM, hTRiC- βTub alone was prepared at 1 µM. All samples were prepared in 50 mM HEPES pH 7.4, 50 mM NaCl and 1 mM DTT. TRiC samples were incubated at 37 °C for 30 min with nucleotide to reach equilibrium/closed state. Fresh DSS crosslinker dissolved in DMSO was added to a final concentration of 1 mM, TRiC samples were incubated at 37 °C for 1 h with crosslinker, hPFD-α/βTub alone was incubated at room temperature for 1 h at room temperature. Crosslinking was quenched by incubation for 30 min after addition of 1 M Tris pH 7.4 to a final concentration of 100 mM. Crosslinked samples were processed as described previously (Leitner et al., 2014). In brief, processing steps included reduction and alkylation of cysteine residues with tris(2-carboxyethyl)phosphine and iodoacetamide, respectively, sequential digestion with endoprotease Lys-C (Wako) and trypsin (Promega), clean-up using solid-phase extraction (Waters Sep-Pak tC18 cartridges) and fractionation of the purified digests by size exclusion chromatography (SEC; GE Superdex Peptide PC 3.2/300).

Liquid chromatography-tandem mass spectrometry (LC-MS/MS) was performed on an Easy nLC-1200 HPLC system coupled to an Orbitrap Fusion Lumos mass spectrometer (both ThermoFisher

Scientific). Peptides were separated by reversed-phase chromatography on an Acclaim PepMap RLSC C18 column (250 mm × 75 µm, ThermoFisher Scientific) at a flow rate of 300 nl/min. The mobile phase gradient was 11 to 40% B in 60 min, with mobile phases A = water/acetonitrile/formic acid (98:2:0.15, v/v/v) and B = acetonitrile/water/formic acid (80:20:0.15, v/v/v). MS/MS data were acquired in the data-dependent acquisition top speed mode with a cycle time of 3 s. Precursor and fragment ion spectra were acquired in the Orbitrap at 120000 and 30000 resolution, respectively. Precursor ions with a charge state of +3 to +7 were fragmented in the linear ion trap at a normalized collision energy of 35%. Dynamic exclusion was enabled for 30 s after one sequencing event.

MS/MS spectra were analyzed using xQuest, version 2.1.5 (available from https://gitlab.ethz.ch/leitner_lab/xquest_xprophet). The database contained the entries of all human TRiC and prefoldin subunits, human tubulin beta (TBB5_HUMAN), and insect tubulins as possible contaminants. No other contaminant proteins were observed at relevant levels. Search parameters for xQuest included: enzyme = trypsin, maximum number of missed cleavages = 2, carbamidomethylation of Cys as fixed modification, oxidation of Met as variable modification, Lys and protein N terminus as cross-linking sites, mass error tolerances of ±15 ppm at the MS1 level and ±20 ppm at the MS2 level. Search results were further filtered with stricter mass tolerances depending on the dataset, and identifications were required to have an xQuest delta score <0.9, a TIC score >0.1, and a minimum of four bond cleavages overall or three consecutive bond cleavages per peptide. A parallel search against the reversed and shuffled sequences of the database entries revealed no decoy hits fulfilling the search and filter criteria, suggesting the false discovery rate is close to 0% for the selected score thresholds. Mapping of XLs was performed using xiVIEW (Graham, M., Combe, C. W., Kolbowski, L. & Rappsilber, J. xiView: A common platform for the downstream analysis of Crosslinking Mass Spectrometry data. *doi: 10.1101/561829)*

**Native Mass Spectrometry**

PFD and PFD-βTub were diluted 20-fold from storage buffer into 250 mM ammonium acetate, pH 7.0, before completing buffer exchange using 10 kDa MWCO Amicon centrifugal filters (Millipore Sigma). Final concentrations were estimated by UV absorbance to be between 10 and 20 µM PFD. Native mass spectra were then recorded within several hours, during which proteins were kept on ice. Samples were injected using borosilicate emitters (ThermoFisher Scientific ES380) into a QExactive Plus EMR orbitrap mass spectrometer (ThermoFisher Scientific). The quadrupole was set to transmit m/z range 1,000 to 10,000. Resolution was 17,500 at 200 m/z for a transient time of 64 ms. Injection time was 50 ms and automatic gain control was off. The capillary voltage was 1.2 kV and S-lens RF was 100. Microscans were grouped sequentially in sets of 10. In-source activation was 50 V and activation in the higher-energy collisional dissociation (HCD) cell was 20 V, based on values found to sharpen peaks without discernable complex dissociation. Pressure in the HCD cell (nitrogen) was 5e-10 mbar. For transmission of PFD, the injection flatapole was set to 20 V, the inter flatapole lens to 12 V, and the bent flatapole to 6 V; for PFD-β-tubulin transmission, these values were respectively 12, 10, and 3 V. Raw data were summed and visualized in XCalibur v4.1.31.9 (ThermoFisher Scientific). Initial deconvolution was performed using UniDec v3.1.0 (Marty et al. 2015). Masses were measured manually by minimizing error over different charge state assignments. Each charge state arising from PFD and PFD-β-tubulin was assigned to the open or compact conformation based on Gaussian peak envelopes. Raw signal intensities of each peak were recorded from XCalibur, summed within assigned conformations, and used to derive fractional occupancies.

**Cryo-EM specimen preparation and data collection**

*Apo-TRiC:* 3 µl of apo-TRiC sample was vitrified at 2 mg/ml was applied to 200-mesh R1.2/1.3 holey-carbon grids (Quantifoil) coated with Poly-L-lysine and vitrified using Vitrobot Mark IV (Thermo Fisher Scientific, CMCI in Seoul). 6084 Movies were collected on a Titan Krios (Thermo Fisher Scientific) equipped with a Falcon 4 (Thermo Fisher Scientific) detector.

*PFD-βTub-TRiC:* 0.5 µM TRiC sample was mixed with 2 µM PFD-βTub complex at RT. CryoEM grids (Quantifoil R1.2/1.3 200 Cu) were glow discharged (PELCO easiGlow) for 45 s. 0.07 % Octyl-beta-glucoside was mixed with the sample prior to vitrification. Sample was vitrified using Vitrobot Mark IV, where 2.7 µl sample was applied on the cryoEM grids and blotted for 3 s. 11,796 movies were collected on a Titan Krios (Thermo Fisher Scientific) equipped with the K2 Summit (Gatan) detector.

*TRiC-βTub with ATP-AlFx*: 1 µM co-purified TRiC-βTubulin sample was incubated with ATP-AlFx for 1 hr at 37 °C. CryoEM grids (Quantifoil R1.2/1.3 200 Cu) were glow discharged (PELCO easiGlow) for 45 s. Sample was vitrified using a Gatan Leica GP plunger, where 2.7 µl sample was applied on the cryoEM grids and blotted from the back for 3~5 s. 31,204 movies were collected on Titan Krios (Thermo Fisher Scientific) equipped with the K2 Summit (Gatan) detector.

**Data processing and 3D refinement**

All image processing was done in RELION 3.1 (Zivanov et al., 2018) and cryoSPARC v3.2. (Punjani et al., 2017). Computing resources were utilized in the $S^2C^2$ SLAC national facility and at CMCI at Seoul National University.

*Apo-TRiC:* Movies were aligned in 5 x 5 patches in MotionCor2, (Zheng et al., 2017) and CTF parameters were estimated with GCTF (Zhang, 2016). Utilizing template-based autopicking in cryoSPARC v3.2, 2,100,259 particles were initially picked. After 2D classification and removing bad particles, 945,248 particles were subjected to 3D heterogeneous refinement using Ab initio model in cryoSPARC. After further 3D classification and CTF refinements, non-uniform refinement was performed using 662,744 particles yielding a 3.11 Å map of apo-TRiC based on the gold-standard Fourier shell correlation (FSC) at 0.143. This analysis workflow is illustrated in Fig. S3.

*PFD-βTub-TRiC:* Movies were aligned in 5 x 5 patches in MotionCor2, and CTF parameters were estimated with GCTF. After initial template-based picking of 1,621,636 particles, 2D classification was performed. After selection, 443,858 particles were used for 3D classification yielding 2 major classes, Prefoldin bound TRiC with 194,013 particles and non-bound TRiC with 249,845 particles. Using non-uniform refinement, prefoldin bound and non-bound TRiC was reconstructed at a resolution of 3.9 Å and 4.2 Å, respectively. This analysis workflow is illustrated in Fig. S3.

*TRiC-βTub with ATP-AlFx:* 21,486 movies were aligned in 5 x 5 patches by MotionCor2 (v), and CTF parameters were estimated with CTFFIND (v 4.1). Particles picked with a template matching method were subject to multiple rounds of 2D classification followed by 3D classification. 819,553 particles in open state were used to reconstruct an open state TRIC-βTub map to 3.8 Å, and 529,181 particles were used to obtain a 2.7 Å C1 symmetry map. This analysis workflow is illustrated in Fig. S4A.

*Compositional heterogeneity analysis of TRiC-βTub with ATP-AlFx:* 529,181 particles of closed state are aligned to the C2 (D1) symmetry axis to obtain a 2.5 Å consensus map of closed state TRIC-tubulin. For compositional analysis of tubulin in the TRiC chamber, the star file of the related particles was extended 2 times by relion_particle_symmetry_expand with D1 symmetry to cover the orientation for each of two rings in each particle. 3D focused classification of the tubulins was performed with a tubulin mask but without symmetry and orientational search. The identified tubulin occupied rings were thus traced back to the original double-ring TRiC particles for the tubulin occupancy analysis. This analysis workflow is illustrated in Fig. S4A.

*Conformational heterogeneity analysis of TRiC-βTub with ATP-AlFx:* The tubulin occupied rings were further subject to focused classification for conformational heterogeneity analysis. In brief, a tubulin mask was applied to the related rings in multiple 3D classifications without orientational search, four tubulin intermediate conformations were identified and each conformation was further locally refined to high resolution. This analysis workflow is illustrated in Fig. S4B.

**Model building**

*Prefoldin-tubulin TRiC:* Model building started from previous prefoldin-TRiC model (PDB: 6NR8). The initial model was fitted into the density map and manually refined in COOT (Emsley et al., 2010). Then, the model was refined on the Namdinator server using MDFF (Kidmose et al., 2019) and Phenix real space refinement default options (Kidmose et al., 2019). After few rounds, the model was further corrected using Phenix and COOT (Afonine et al., 2018).

*Copurified tubulin TRiC under ATP-AlFx condition:* The model of TRiC in the closed form (PDB ID: 7LUM) reference was rigidly fit into the closed state TRiC density by rigid body fitting with Fit in Map tool from Chimera v1.14. This fitted model was further refined with phenix.real_space_refine in Phenix v1.18.1, ISOLDE v1.1.0. And refined models were inspected and adjusted in COOT. The model of tubulin (PDB: 6I2I) was rigidly fitted to each tubulin intermediate density, and manually adjusted by COOT and ISOLDE v1.1.0. All adjusted models were then refined using phenix.real_space_refine in Phenix.

All models were validated by Q-score (Pintilie et al., 2020) and phenix.validation_cryoem (Afonine et al., 2018). Difference maps showing the nucleotide density in TRiC-tubulin closed state were calculated between the complex density map and the map calculated from the model of protein only, generated by phenix.real_diff_map. The figures of the difference map were generated by Chimera with the same contour level 3 sigma. All other figures were generated by Chimera and ChimeraX.

**PISA analysis**

The interactions between TRiC and tubulin model in each state were calculated using the PISA server (https://www.ebi.ac.uk/pdbe/pisa/).

**Logo Plot generation**

Residue conservation was compared across 393 mainly eukaryotic with some archaeal species (group II chaperonins) for residues predicted to make interactions by PISA. Plots were generated using *EDlogo* plot (Dey et al., 2018).

**Sequence alignment and phylogenetic tree building**

Multiple sequence alignments for tubulin and tubulin homologues were generated using the multiple sequence alignment tool T-coffee Espresso and visualized (Di Tommaso et al., 2011). A phylogenetic tree of tubulin and tubulin homologs was made in MEGA X software (Hall, 2013).

#### DATA AVAILABILITY

The 3D cryoEM density maps have been deposited in the Electron Microscopy Data Bank under the accession number EMD: EMD-32822, 32823, 26089, 26120, 26123, 26131. Coordinates have been deposited in the Protein Data Bank under the accession number PDB: 7WU7, 7TRG, 7TTN, 7TTT, 7TUB. The mass spectrometry proteomics data have been deposited to the ProteomeXchange Consortium via the PRIDE partner repository with the dataset identifier PXD030590.
